# Tumor microenvironment signature associated with morphology in systemic ALK-positive anaplastic large cell lymphoma

**DOI:** 10.64898/2026.01.20.700117

**Authors:** Chloé Bessiere, Loélia Babin, Sandra Dailhau, Philippe Gaulard, Stéphane Pyronnet, Fabienne Meggetto, Laurence Lamant

## Abstract

Peripheral T-cell lymphomas (PTCL) are heterogeneous entities whose tumoral microenvironment (TME) may influence disease phenotype and outcome. To dissect their immune and stromal composition, we used two complementary algorithms, CIBERSORTx and MCP-counter, on Affymetrix data from 255 patients whole-tissue biopsies, including 78 systemic anaplastic large cell lymphomas (ALCL). Clustering based on inferred cell proportions revealed a clear separation between ALCL and other PTCL subtypes, including angioimmunoblastic T-cell lymphoma (AITL) and PTCL-not otherwise specified (PTCL-NOS). ALCL-enriched clusters were characterised by granulocytes and macrophages lineages, mast cells, NK cells, memory CD4+ T-cells, as well as fibroblast and endothelial signatures, whereas as expected, AITL cluster was enriched for T-follicular helper cells, B-cells and macrophages. Finally, we focused on 66 ALCL cases with relapse risk and morphological subtype clinical annotations. Distinct TME features enriched in M1 macrophages and monocytes were associated with adverse outcomes and non-common morphological variants.

Transcriptomic analyses of mononuclear phagocytes across Affymetrix and RNAseq datasets confirmed distinct clustering according to cell morphology. These analyses identified potential biomarkers associated with uncommon variants and absent from ALK+ ALCL cell lines, confirming their TME origin. Macrophage- and monocyte-related signatures emerged as key contributors in ALK+ ALCL patients heterogeneity, linking TME patterns to tumor morphology and prognosis. These signatures may serve as biomarkers for patient risk stratification and guide the development of targeted therapies.

## Introduction

Peripheral T-cell lymphomas (PTCLs) represent a biologically heterogeneous group of non-Hodgkin lymphomas arising from mature T lymphocytes. They most commonly present as nodal disease and include several subtypes with distinct clinical and pathological characteristics, such as systemic anaplastic large cell lymphoma (ALCL), angioimmunoblastic T-cell lymphoma (AITL), and PTCL-not otherwise specified (PTCL-NOS).

ALCLs are mostly CD4+ phenotype, but can sometimes be CD8+, while still strongly expressing the lymphocyte activation marker CD30 (1). These high-grade non-Hodgkin lymphomas (NHL) account for 15% of paediatric and 1-2% of adult NHL. The majority of paediatric cases and around half of adult cases (2,3) are associated with a translocation involving the *ALK* gene. In ALK-positive (ALK+) ALCLs, the ALK fusion protein promotes oncogenicity by activating several signalling pathways, including PI-3K, MAPK and JAK/STAT pathways (4).

These tumors are relatively chemosensitive, showing high response rates to first-line chemotherapy regimens. However, despite different treatment protocols and durations, 2-year event-free survival (EFS) remains between 65 and 75%, and overall survival (OS) between 70 and 90% (5,6). Several factors have been associated with poorer prognosis in the paediatric population including the small cell (SC) and lympho-histiocytic (LH) morphological variants (1), the detection of early minimal residual disease in blood and/or bone marrow (7), and a low anti-ALK serum antibodies titer (8,9). For refractory or relapsed ALK+ ALCLs, new treatment options have recently emerged, such as brentuximab vedotin, an antibody–drug conjugate targeting the CD30 antigen on the surface of ALCL cells (10), or ALK inhibitors (11). Immune checkpoint inhibitors have also shown promise in solid cancers, although their use in hematologic neoplasms, particularly T-cell NHL, remains less documented. PD-L1 expression is frequently observed in ALK+ ALCLs through STAT3-mediated pathways (12,13) providing a rationale for targeting immune checkpoints (14,15).

Beyond these targets expressed by tumor cells, the tumor microenvironment (TME) plays a key role in cancer immunoediting by regulating tumor growth, dissemination, and therapeutic resistance (16–18). Furthermore, the cellular and molecular constituents of the TME could reveal additional immunotherapeutic targets. This study aimed to decipher the diversity of immune infiltrating cells at diagnosis, first comparing ALCL with other PTCLs, and then focusing on ALK+ ALCL before treatment, to identify TME relevant signatures associated with morphological and/or clinical features.

## Materials and Methods

### ALCL cohorts and clinical data

78 patients diagnosed with a systemic ALCL (61 ALK+, 17 ALK-), 83 with AITL, 71 with a PTCL-NOS and 23 undetermined AITL/PTCL-NOS were retrieved from the translational T-cell lymphoma research consortium (TENOMIC) of the Lymphoma Study Association (LYSA) and PAIR lymphoma project. All the patient biopsies were obtained at diagnosis and cases had been reviewed by at least two expert hematopathologists, according to the criteria of the 2017 WHO classification (19). The percentage of tumor cells was assessed using CD30 staining in ALCL, combined with morphology in other PTCLs and accounted for 30 to 90% of the lymph node involvement (20,21).

Gene probes intensity levels using HG-U133 plus 2.0 Affymetrix GeneChip arrays (Affymetrix, Santa Clara, CA) were previously available for all these fresh-frozen tumors (20–23). Among the cohort of 61 systemic ALK+ ALCL **(Table S1)**, 48 tumor samples were already used to identify a molecular signature based on microarray data that was associated with clinical outcome (26 with relapse/progression versus 22 non-relapse) and 39/48 (18 relapsing and 21 non-relapsing) were further retained for RNAseq (full RNA 150-bp paired-end sequencing after ribodepletion) (23). The study was approved by the ethics committee of the TENOMIC program (Comité de Protection des Personnes Ile-de-France IX 08-009).

### ALCL cell lines and patient-derived xenograft (PDX)

ALK+ ALCL cell lines KARPAS-299, SU-DHL-1, SUPM2, Pio and COST, carrying the t(2;5)(p23;q35) translocation were obtained from DSMZ (German Collection of Microorganisms and Cell Culture) or established in our laboratory (24). NPM1-ALK transformed and immortalized T-cell models M1 (lentiviral NPM1-ALK+) (25) and ALKIma1 (CRISPR-Cas9 induced NPM1-ALK+) (26) were produced in our laboratory. All ALK+ ALCL cell lines were cultured in RPMI-1640 medium (Invitrogen #61870044) supplemented with 20% FBS (PAN BIOTECH #500105M1M). Normal CD3+ lymphocytes activated (n=3) or non-activated (n=3) are paired and were obtained from 3 different donors (25). The DN03 and SG17-PDX were provided by G. Inghirami (27).

### Data preprocessing

Affymetrix probe intensities data were processed using the *affy* R package and normalized with the *rma* function by a robust multi-array average (RMA) expression measure (28). We then used the collapse microarray tool (https://sites.google.com/site/fredsoftwares/products/collapse-microarray) to get a single normalized expression value by gene, using the provided Affymetrix U133+2 corresponding to our samples chip reference.

The RNAseq raw FASTQ files were directly processed with Kallisto v0.46.2 pseudo-alignment method (29) to quantify transcript abundances. It was performed with a 100 bootstrap value (-b 100) and with the *--rf-stranded* parameter. As a reference annotation, we used a transcriptome index constructed from the Ensembl project’s transcriptome v108 (cDNA + ncRNA) from which we excluded alternative loci scaffolds. We generated gene counts scaled using the average transcript length, averaged over samples and to library size with the *tximport* R package and *countsFromAbundance* argument equal to “lengthScaledTPM”.

### Inference of TME cells into ALCL patient samples

To infer the proportions of immune and stromal cells in the PTCL samples, we used the CIBERSORTx algorithm based on deconvolution (30) and the MCP-counter (microenvironment cell populations-counter) based on robust transcriptomic markers expressed mainly in one single cell subtype (31). The CIBERSORTx was used with the included precomputed LM22 gene signature matrix (547 genes) to infer 22 human immune cell subtypes (B-cells, T-cells, natural killer cells, macrophages, DCs, and myeloid subtypes) into bulk tumor samples. Affymetrix and RNAseq normalized expression tables were uploaded on the CIBERSORTx web portal (https://cibersortx.stanford.edu/runcibersortx.php) and we run the Impute Cell Fractions module with 500 permutations enabling batch correction (B-mode). We also applied the MCP-counter method to predict, for each of our samples, the absolute abundance of 10 human immune and stromal cell subtypes. The 8 immune cell types serve as a control for CIBERSORTx estimation and the 2 stromal cell populations are used to enrich included cell types in our study. To have uniform and comparable values of cell type estimations between the 2 tools, we reduced and centered all the estimation scores (Z-scores). Whereas CIBERSORTx relies on deconvolution, and therefore does not allow each discriminating gene to be directly associated with cell subtypes, MCP-counter uses a list of robust marker genes by cell type. We indeed used these marker gene lists to interpret the differences between ALCL and other PTCLs. To compare the different estimated cell types between patient groups, we applied a Wilcoxon statistical test and corrected the p-value with the Hochberg method.

### Clustering for TME-infiltrating cells

To compute the unsupervised clustering of samples based on cell-type scores, we used the *ConsensusClusterPlus* R package (32). This method provides quantitative and graphical information to determine the best number of clusters (k). Here we selected 90% samples resampling (pItem) taking all cell-types (100% for pFeature). We also defined a maximum k of 4 and a 500 times resampling (reps) to ensure classification stability. We also used hierarchical agglomerative clustering (clusterAlg=“hc”) based on Euclidean distance (distance=“euclidean”) and Ward’s linkage (innerLinkage=“ward.D”) for the grouping.

### Pathway gene signatures and Sample Enrichment Score (SES)

The gene signatures related to macrophage and monocyte pathways were extracted from the WikiPathways and Gene Ontology Consortium databases. These signatures are listed into the **Tables S2 and S3** for macrophages and monocytes respectively.

To evaluate the enrichment of a gene signature in our samples, we applied the open source software AutoCompare_SES (https://sites.google.com/site/fredsoftwares/products/autocompare_ses) (33). We used a “greater” distribution side to extract enriched gene sets. The SES ranges from 0 to 100. We calculated these scores in both our ALCL patient Affymetrix and RNAseq datasets from which we have the morphology information, and we evaluated the significativity of a signature by comparing common-type (CT) and non common-type (nonCT) patient groups with a wilcoxon test using the *stat_compare_means* R function. If the p-value is <0.05 in both the Affymetrix and RNAseq datasets, we keep the signature as relevant to discriminate between our two patient groups. In the heatmaps, we represented the SES normalized scores with a scaled normalization (reduced and centered Z-scores).

### Transcriptomic signature associated with prognosis clusters

For each retained macrophage or monocyte signature, we have a list of genes. To perform a transcriptomic analysis, we keep the union of the different gene lists for the macrophages on one side and monocytes on the other side. In the end, we keep only, from these 2 union lists, the genes that are also differentially expressed in the differential expression analysis (DEA) from lengthScaledTPM kallisto counts. The DEA was done with Sleuth on the 38 ALCL patient RNAseq gene counts from kallisto, between the CT (n=22) and nonCT (n=16) conditions. We defined a threshold of more than 1 for the abs(logFC) and a false discovery rate adjusted p-value (Benjamini-Hochberg) < 0.05. We also did a DEA with the *limma* R package, based on Affymetrix RMA normalized gene expression and using the same corrected p-value threshold of 0.05 with an abs(logFC) > 1, to define the most confident genes among the signatures. Gene lists for both RNAseq and Affymetrix are in **Table S4.**

### K-mer based gene expression data

To verify the specific microenvironment expression of the selected mononuclear phagocyte (macrophages and monocytes) genes, we checked for their expression in both activated or not CD3 T-lymphocytes and ALCL ALK+ cell lines or PDX. To have a precise and quick expression profile of these genes in all sub-cited samples, we used a recent k-mer based approach. Briefly, we generated for each gene all the specific subsequences of 31 nt called k-mers from their canonical sequence using the Kmerator tool with the ensembl reference transcriptome v108 (-r 108) (34). We then used transipedia.org (35) to retrieve in a fraction of second the mean expression value by gene, in all samples previously indexed with Reindeer (36) and in private access, and normalized by billion of k-mers.

## Results

### Microenvironmental cell signatures delineate ALCL among PTCL

To profile the TME components in ALCL and two other types of PTCL (AITL and PTCL-NOS), a computational approach was applied to Affymetrix gene expression data obtained at diagnosis from tissue biopsies of 255 PTCL including 78 ALCL samples. Two algorithms, CIBERSORTx and MCP-counter (30,31), were used to characterise the proportions of 22 immune cell subtypes and 2 stromal cell subtypes *in silico*. To ensure robust classification, the *ConsensusClusterPlus* R package was applied, providing quantitative evidence for the optimal number of clusters and samples assignment. Clustering revealed a clear separation between ALCL and other PTCL subtypes **(Figure 1, Figure S1A)**, driven by their immune and stromal landscapes. Indeed, analysis of the TME in ALCL and non-ALCL groups **(Figure 1)** showed that the most homogenous cluster 1 is significatively enriched in ALK+ and ALK− ALCL, suggesting that TME signatures may be clinically relevant for trials using TME-targeted therapies. Similar results were obtained in the analysis of ALCL and AITL **(Figure S1A)** with two major clusters enriched respectively with ALCL and AITL samples. Among the ALK+ ALCL enriched cluster microenvironment, the most prominent immune cell subsets were monocytes (*p*val = 6.8e-12), mast cells resting (*pval* = 1.9e-16), M2-like macrophages (*pval* = 8.9e-11), natural killer (NK) resting cells (*pval* = 1.4e-22), memory CD4+ activated cells (*pval* = 1.6e-20) and neutrophils (*pval* = 5.5e-14) **(Figure 1, Figure S1)**. Unlike CIBERSORTx based on the principle of deconvolution, MCP-counter relies on a set of well-chosen transcriptomic marker genes for 10 different cell types. CSF1R, PLA2G7, RASSF4, FPR3, ADAP2, KYNU and TFEC were significantly enriched from the MCP-counter monocytic lineage (which included macrophages) in ALCL **(Table S5)**. In addition, analysis revealed that fibroblasts and endothelial signatures were more prevalent in ALCL compared with other PTCL **(Figure 1, Figure S1A)**. Moreover, AITL samples were characterised by the overexpression of genes associated with macrophages M0 cells, γδ T-cells and T follicular helper (TFH) cells **(Figure S1B)**. As AITL originates from TFH cells and is usually associated with significant TME (37), these results further support our approach to estimate immune cell composition. However, given the potential overlap between transcriptional programs of memory CD4+ activated cells or CD8+ T-cells and tumoral ALCL cells, as previously reported for B-cell signatures in B-cell malignancies (38), memory CD4+ activated and CD8+ cells were excluded from subsequent analyses.

**Figure 1:**
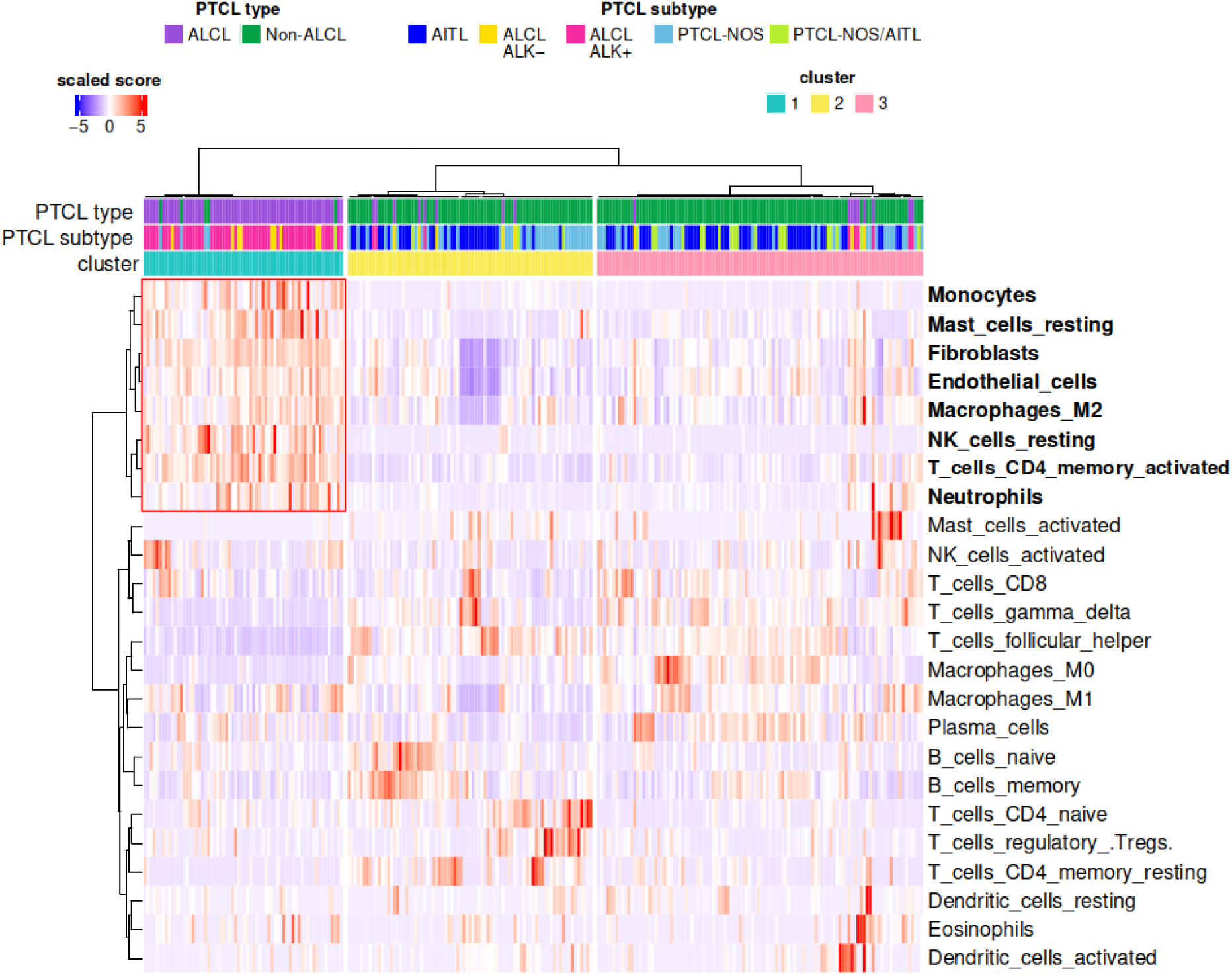
microenvironmental cell signatures of ALCL versus other PTCL. The heatmap represents the Z-score (normalized proportion of cell types) of the 22 different immune cell types from CibersortX and 2 additional stromal cell types from MCP-Counter. 255 samples are included : 78 ALCL (61 ALK+ and 17 ALK− ALCL), 177 non-ALCL PTCL including 83 AITL, 71 PTCL-NOS and 23 undetermined AITL or PTCL-NOS. The red frame highlights the cell subtypes enriched in ALCL. PTCL = Peripheral T-Cell Lymphoma; ALCL = Anaplastic Large Cell Lymphoma; AITL = Angioimmunoblastic T-Cell Lymphoma; NOS = not otherwise specified.

### The tumor microenvironment signature is associated with morphological variants in ALK+ ALCL

To investigate whether the immune cell signatures of the TME are important for predicting outcomes, a clustering analysis was performed on a subgroup of 66 ALCL with clinical information about early relapse after chemotherapy and/or with known morphological subtype. This approach was based on earlier findings that nonCT morphology is significantly and independently associated with relapse or early progression (1). As previously noted, the memory CD4+ activated cells and CD8 T-cell signatures were excluded from this *in silico* analysis due to the potential transcriptional overlap between normal infiltrating lymphoid cells and ALK+ lymphoma T-cells. Unsupervised analysis of the 66 ALCL patients from their immune and stromal profiles identified two distinct clusters: tumors from non-relapsing patients are enriched in cluster 1, while cluster 2 is predominantly composed of early-relapsing cases. To confirm the link between TME composition and relapse risk, the estimated proportion of each immune cell type and their associated score were compared between relapsing and non-relapsing tumors. No statistically significant differences were found in immune cell composition between the two groups. By contrast, when ALK+ ALCL with CT morphology (the most frequently observed) were compared with nonCT morphological variants, already shown to be of prognostic value (1), a clearer clustering pattern emerged **(Figure 2A)** with a cluster 2 composed in majority of nonCT variant tumors. This stratification is linked to significant differences in 3 immune cell populations: M1 macrophages, monocytes and dendritic cells activated, with enrichment of M1 macrophages and monocytes in the nonCT morphological variants **(Figure 2B)**. In the following analysis, dendritic cells were not studied, as their scores are very low (<1%) in the majority of samples **(Figure S2)**.

**Figure 2:**
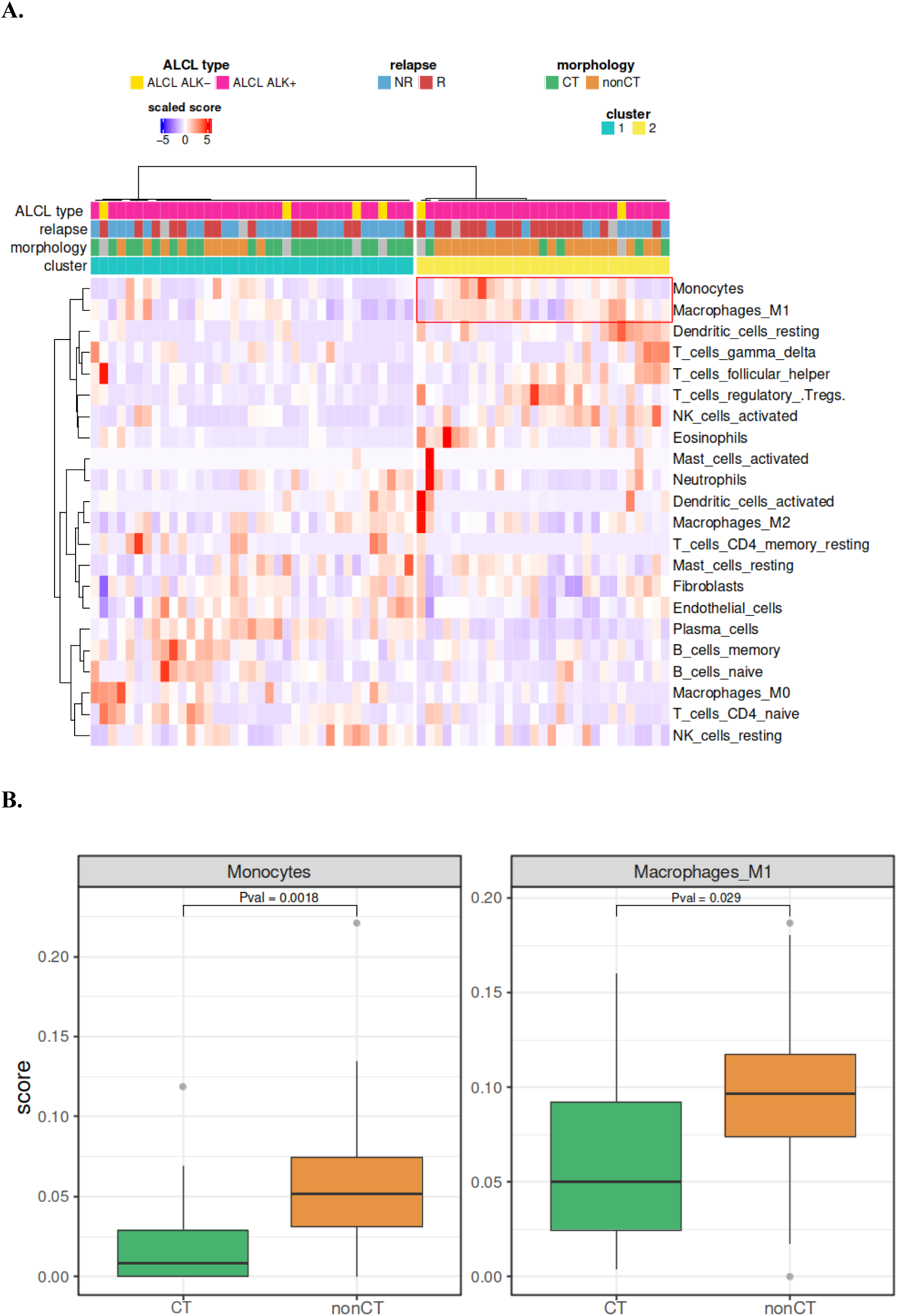
TME exploration in ALCL patients with their associated clinical data : ALCL type, early relapse and morphology. **(A)** The heatmap represents the Z-score (normalized proportion of cell types) of 20 different immune cell types from Cibersort and 2 additional stromal cell types from MCP-counter. 66 ALCL patient samples with at least one information on relapse or morphology are included. The red frame highlights the cell subtypes significatively enriched in nonCT patients. **(B)** Monocytes and Macrophages M1 are both enriched into nonCT patients compared to CT ones. CT and nonCT variants groups include 28 and 32 patients respectively. The p-values were calculated with a Wilcoxon test corrected by the Hochberg method. ALCL = Anaplastic Large Cell Lymphoma; NR = non-relapsing; R = relapsing, CT = Common-Type variant; nonCT = non-Common-Type variant. In boxplots the center line represents the median, boxes indicate the interquartile range, the whiskers show the 1.5 interquartile range.

### The macrophage imprint is reflected in the tumor morphology

To evaluate the predictive value of macrophage-related transcriptional activity for morphological classification, publicly available databases of macrophage gene signatures were first explored. Twenty-five gene sets related to macrophage pathways were identified **(Table S2)**. For each signature, a sample enrichment score (SES) was computed across 60 tumor samples with well-defined morphological subtypes **(Figure S3A)**. Of these, 17 showed significant differential enrichment between common and non-common morphological variants in the Affymetrix dataset and their expression pattern gave rise to three distinct clusters. Cluster 1 was enriched in cases with CT variants, while cluster 2 predominantly included nonCT variants **(Figure S4A)**. To validate these findings using a transcriptomic dataset generated by another technique, the same analytical pipeline was applied to an RNAseq dataset that had previously been generated from a subset of 38 patients (23) **(Figure S3A)**. This analysis identified three clusters, with cluster 1 enriched in CT morphological variants and cluster 2 enriched in nonCT variants. This confirmed the robustness of the signature **(Figure S4B)**. Focusing on signatures that were significant in both datasets, six robust macrophage-related gene sets were identified **(Figure 3, Figure S3B)**. These included pathways involved in the activation of macrophages during immune responses, as well as their differentiation and proliferation. As shown in **Figure 3A** and **Figure S4**, the enrichment patterns of these 6 gene signatures define 3 distinct clusters : (1) a cluster with low overall macrophage activity, primarily comprising tumors with a common morphology; (2) a cluster with high enrichment scores, predominantly composed of nonCT variants; and (3) a heterogeneous intermediate group containing tumors of both morphologies. Whatever the Affymetrix or RNAseq data set used, ALCL samples (35 on 38 common patients between RNAseq and Affymetrix samples) tested with both methods are roughly classified in the same group. Among the 13 CT samples from the RNAseq low macrophage activity signature **(Figure S4B, cluster 1)** only one was assigned to the intermediate group in the Affymetrix dataset (marked as underlined). Similarly, of the 7 patients with high macrophage activity according to RNAseq **(Figure S4A, cluster 2)**, only one was differently classified in the Affymetrix data. These results highlight the robustness of macrophage-related transcriptional patterns across different platforms, as well as their correlation with morphological subtypes. Next, a shortlist of transcriptomic biomarkers associated with morphology was compiled. Using the 89 genes included in the six macrophage-related signatures **(Figure 3A, Figure S3B, Table S6)**, differential expression analysis was performed between tumors with common and non-common morphology (*pval* < 0.05 and abs(logFC) > 1). Seven genes (IL34, IFNG, LYZ, CSF2, miR181b1, LBP and ZBTB46) emerged as differentially expressed in the RNAseq dataset **(Figure 3B, Figure S3B)**. To confirm that these genes were associated with the TME rather than with tumor cells, their expression was assessed in a panel of ALK+ ALCL cell lines and two patient-derived xenografts (PDX). With the exception of ZBTB46, which was consistently expressed in tumor cells, the other six genes showed absent or very low expression in ALK+ ALCL models **(Figure S5A)**, supporting the hypothesis that their differential expression in patient samples reflects the infiltration or activation of cells from the macrophagic lineage. The constitutive expression of ZBTB46 in ALK+ cell lines has previously been reported (39,40) and is consistent with our observations, therefore this gene was excluded from the list of macrophage-associated TME biomarkers. Among the 6 remaining genes, IFNG, LYZ, and LBP were also validated in the Affymetrix dataset, strengthening their potential as macrophage-related markers **(Figure S5B)**.

**Figure 3:**
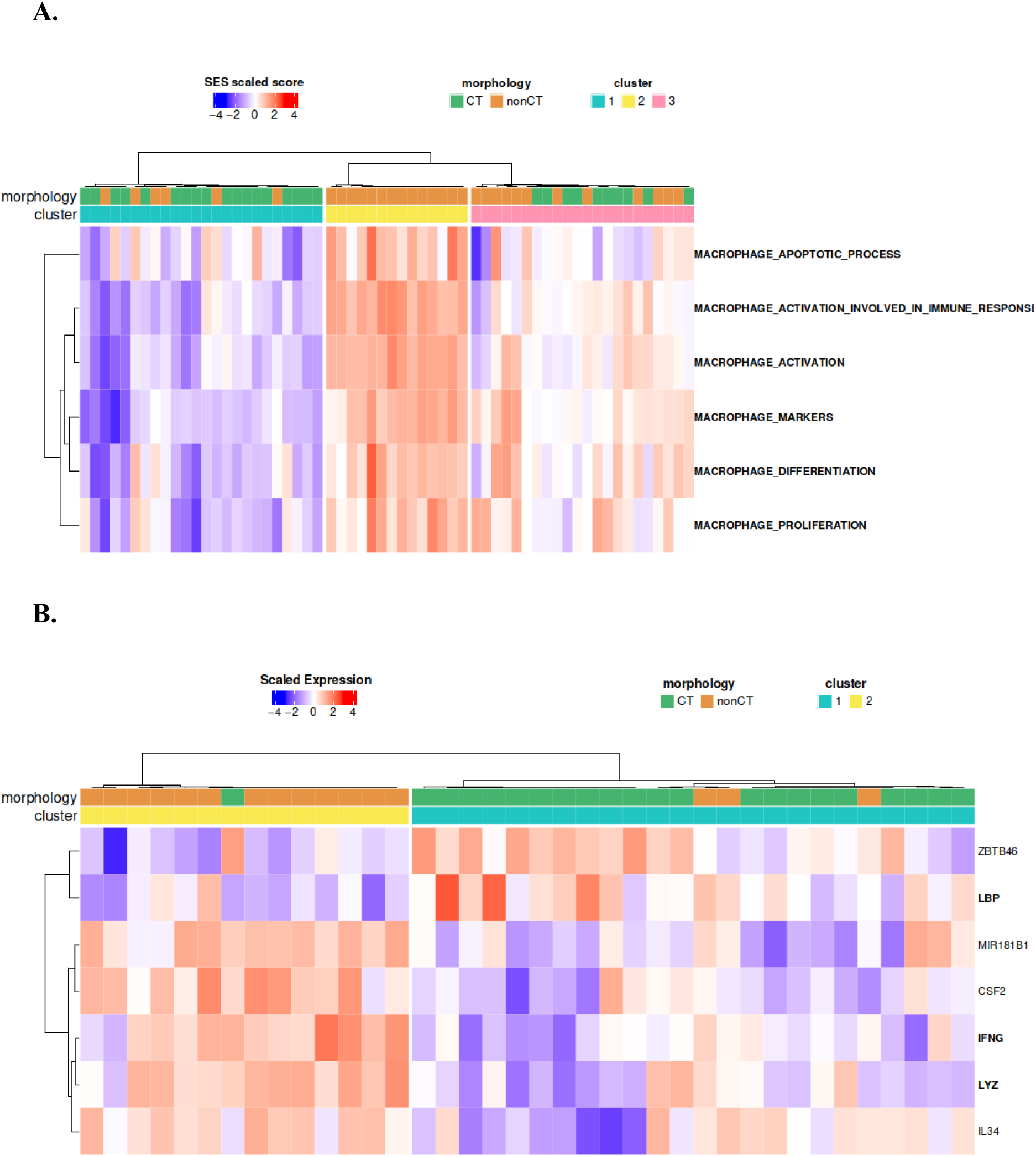
Macrophage signature of ALCL tumors. **(A)** normalized SES scores from the significatively differential gene signatures between common and non-common type variants in both the Affymetrix and RNAseq datasets. Affymetrix SES scores are represented. **(B)** best transcriptomic signature to discriminate between common and non-common type variants using RNAseq data.

It is well established that certain morphological variants of ALK+ ALCL, namely the LH subtype, are enriched in histiocytes. However, in the group of 14 nonCT cases **(Cluster 2**, **Figure 3A)**, only 7 cases are LH ALCL, defined by an obvious histiocytic infiltration on Hematoxylin and eosin staining. The remaining cases consisted of SC variants or composite ALCL, in which areas of the common type were associated with SC regions. Our study suggests that SC variants appear to be more enriched with histiocytes than expected based on routine sections **(Figure S6)**.

### The monocyte imprint is reflected in the tumor morphology

As for macrophages, the ability of monocyte-related transcriptional signatures to predict morphological subtypes was assessed. Using public databases of general monocyte signatures, nine gene sets associated with monocyte pathways were identified **(Table S3)**. For each signature, a SES was calculated in all 60 tumor samples morphologically well-characterized **(Figure S3A)**. Of these, eight showed significant differential enrichment between CT and nonCT morphological variants in the Affymetrix dataset and their expression pattern gave rise to two distinct clusters : cluster 1 was enriched in cases with CT variants and cluster 2 in cases with nonCT variants **(Figure S7A)**. In the RNAseq transcriptomic dataset including 38 patients **(Figure S3A, Table S1)**, two distinct clusters were also identified. Six signatures were also significantly enriched **(Figure S7B)**, resulting in a robust subset of six monocyte gene signatures **(Figure 4A)**, including pathways related to monocyte activation, chemotaxis and extravasation. As seen on **Figure 4A**, the enrichment pattern of these six gene signatures define two clusters : a first one with high enrichment scores, predominantly composed of nonCT variants, and a second one with low overall monocyte activity comprising a majority of tumors with a common morphology. Whatever the Affymetrix or RNAseq data set used, ALCL samples tested with both methods are classified in the same group **(Figure S7, samples in bold)**. Similar to macrophage signatures, the next step was to define a minimal set of monocyte transcriptomic biomarkers associated with morphology. From the 112 genes included in the six monocyte-related signatures **(Table S7)**, differential expression analysis of RNAseq data was performed comparing tumors with CT *versus* nonCT morphology (*pval* < 0.05 and abs(logFC) > 1). This analysis identified seven genes, CLEC7A, CXCL10, CCL19, JAML, MT1G, CCL18, and CCN3 **(Figure 4B)**, as significantly differentially expressed. To validate the association of these genes with the TME rather than with tumor cells themselves, their expression levels were assessed in the panel of ALK+ cell lines. Consistent with an origin in the TME, these genes exhibited undetectable expression in the cell lines **(Figure S8A)**. This observation supports the hypothesis that their differential expression in patient samples reflects the infiltration or activation of cells from the monocytic lineage. Among these, CLEC7A, CXCL10, and CCL19 were also validated in the Affymetrix dataset and were upregulated in nonCT variants **(Figure S8B)**. CCL18 was also validated and found to be downregulated in nonCT variants.

**Figure 4:**
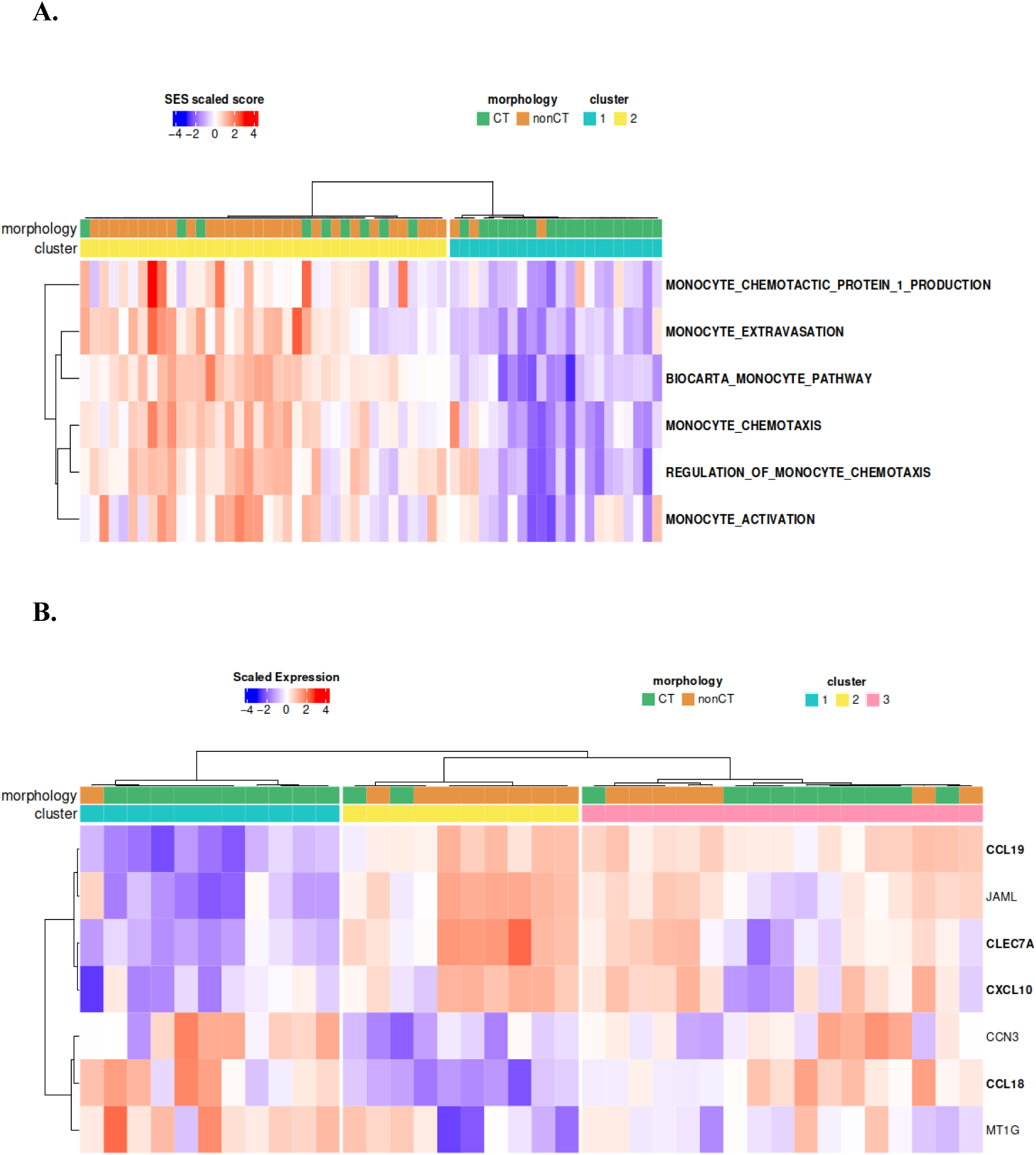
Monocytes signature of ALCL ALK+ tumors. **(A)** normalized SES scores from the significatively differential gene signatures related to monocytes between common and non-common-type variants in both the Affymetrix and RNAseq datasets. Affymetrix SES scores are only represented. **(B)** best monocyte transcriptomic signature to discriminate between common and non-common-type variants based on RNAseq data.

## Discussion

Only a few studies have explored the TME landscape in T-cell lymphomas, mainly in AITL cases (41–43). Until now, due to the rarity of systemic ALCL cases, collecting fresh samples for comprehensive TME analysis using single-cell transcriptomic has been challenging. In this study, already available tumor bulk transcriptomic data were used to explore the TME in ALCL.

Our results show that microenvironmental immune signatures clearly delineate ALCL as a distinct PTCL subgroup. After exclusion of transcriptional programs of memory CD4+ activated cells to avoid overlap between immune CD4+ activated cells and tumoral ALCL cells, the dissection of systemic ALCL TME revealed an enrichment in monocytes, M2-like macrophages, neutrophils, NK resting cells, fibroblasts and endothelial cells. Consistently, Huang et al recently observed a « mesenchymal » main subtype in ALK+ ALCL enriched in M2 macrophages, stromal cells and extracellular matrix (ECM), and a « depleted » subtype characterized by a lower expression of all TME signatures compared to other PTCL subtypes (43).

These immune and stromal cells could participate in a pro-tumoral environment (44), as already reported in Hodgkin lymphoma and DLBCL (45–47). Although the M1/M2 dichotomy is unsatisfying to describe the spectrum of macrophage phenotypes within the TME (48), several genes from the MCP-counter macrophage signature enriched in our ALCLs, FPR3 (Formyl peptide receptor 3), TFEC (a macrophage-specific transcription factor) and CSF1R, polarize macrophages toward an immunosuppressive phenotype termed alternatively activated or M2-like macrophages (49,50). Interestingly, *CSF1R* gene, highly expressed in mononuclear phagocytes, is also involved in macrophage chemo-attraction in solid cancers (51–53). Early trials have evaluated CSF1R inhibitors (54) in lymphomas, suggesting that CSF1R could be a potential druggable target for controlling tumor progression in ALCL, possibly in combination with an anti-PD-1 by depleting and/or reprogramming macrophages.

The CSF1-CSF1R axis is also involved in the crosstalk between macrophages and fibroblasts and more particularly, cancer-associated fibroblasts (CAF), known to facilitate cancer progression through various mechanisms (55). Yet, upon comparison of ALCL with other PTCLs, a fibroblast signature was identified as predominant in ALCL. This could be explained by the TH2 (T helper type 2) profile of ALCL tumor cells as TH2-polarized responses are important for development of fibrosis (56). A “stromal signature” in ALCL associated with a favorable outcome had previously been reported (23), while some ALK+ ALCL cases are classified within the «mesenchymal» subtype (43). These observations suggest that, in ALCL, a balance toward M2 macrophages may promote ECM deposition, with potential downstream consequences on therapeutic response (57). The significant enrichment of mononuclear phagocyte signatures in nonCT ALCL variants is confirmed by our study, and validates the bulk transcriptomic approach employed. Indeed, it is established that LH subtypes are enriched in histiocytes. However, we found enrichment in other morphological variants : SC variants and composite ALCL, where common-type areas coexist with SC regions. This observation supports our hypothesis that SC and LH variants share similar patterns (58), as a substantial proportion of SC is observed in LH variants, and conversely, SC variants show more histiocytes than expected based on routine histological sections. Finally, despite a currently simplistic classification of macrophages into M1 (anti-tumoral) and M2 (pro-tumoral), that should be considered as a spectrum with evolving activation states, it is not surprising that M1 (IFNG, LYZ, CSF2) and M2 (IL34, miR181b1 and LBP) markers are observed in the macrophage signature of ALCL cases explored. Beyond mere enrichment of mononuclear phagocytes in nonCT variants, an overrepresentation of activation and proliferation pathways was revealed.

Zhang et al (59) showed that abundant TAM CD68+ could predict PTCL-NOS of poor prognosis, though, CD68 is a pan-macrophage marker and antibodies such as CD206 or CD163 are needed to better characterize macrophages. Zhu et al also revealed an immunosuppressive TME in relapsed/refractory AITL using single-cell RNAseq (41). Although no significant association was found between immune cell types enumeration and outcome after chemotherapy, these results need further exploration in larger patient cohorts. Given several reports of response to PD-1 blockade in relapsed/refractory ALCL (14,15), it could also be of interest to investigate TME remodeling following such therapy in ALCL. Furthermore, immune cells abundance alone might be insufficient to explain treatment response as spatial interactions between cells could play a critical role (60).

## Conclusion

Our study demonstrates that ALCLs exhibit a distinct microenvironmental immune signature, clearly different from other PTCLs, characterized by an enrichment in monocytes, M2-like macrophages, neutrophils, NK resting cells, fibroblasts and endothelial cells. This TME likely contributes to a pro-tumoral environment. Although these immune and stromal components do not segregate ALCL cases according to clinical behaviour after chemotherapy or morphological subtype, a significant enrichment of mononuclear phagocyte signatures was observed in non-common variants, which are known to have adverse prognostic impact. A deeper understanding of the ALCL-specific TME and its remodeling following immunotherapy, potentially through single-cell RNAseq analysis from more easily available FFPE samples, could help to develop alternative treatments and find predictive biomarkers.

## Acknowledgments

The study was supported by INSERM, Labex Toucan, Association Eva pour la vie, Féderation Grandir sans Cancer, RACCE Lions Club de Lourdes, Association le sourire de Lucie, le marcheur au grand coeur/Fondation les batisseurs d’étoiles, the French Ministry of Health and the French National Cancer lnstitute (CircOma, PRT-K 2022-184), Fondation ARC et Fondation pour la Recherche Médicale supported by its sponsor Nagui and his production company, Banijay Production (EQU202403018019). C.B was supported by Fondation de France, S.D. by Ministère de la Santé and Institut National du Cancer (CircOma, PRT-K 2022-184) and Fonds AMGEN pour la Science et l’Humain. L.B. was supported by Fondation de France, Fondation l’Oréal for Women in Science, Prolific Graine de Chercheur, Association Ellye, Fondation ARC, and Fondation Toulouse Cancer Santé. Scheme for figure S3 was created with Biorender (www.biorender.com).

## Author contributions

C.B., L.L, L.B. and F.M. designed the research study. C.B., L.L. and F.M. analyzed the data. C.B. performed the bioinformatic analysis. L.L. and P.G. were involved in the diagnosis of PTCL. L.L., C.B. and F.M. wrote the paper. S.P. and S.D helped with discussions and critical reading. All the authors have read and agreed to the published version of the manuscript.

## Data availability

Affymetrix and RNAseq datasets from patients with PTCL, including ALK+ ALCL, were previously published (23). All other data generated or analysed during this study are available within the article and its supplementary information files. Additional data or materials are available from the corresponding authors upon request : Laurence Lamant and Fabienne Meggetto.

