## Supplementary material for "Tumor microenvironment signature associated with morphology in systemic ALK-positive anaplastic large cell lymphoma": Supp. methods, tables and figures

**This PDF file includes:**

Supplementary methods

Supplementary tables and Supplementary Figures S1 to S7

### Supplementary Methods

#### ALCL patient informations

Patients provided informed consent, in accordance with the Declaration of Helsinki. ALCL tumor samples were registered at the ‘CRB cancer des Hôpitaux de Toulouse’. According to the French law, CRB Cancer collection has been declared to the Ministry of Higher Education and Research (DC 2020-4074) and obtained a transfer agreement (AC-2020-4031) after approbation by ethical committees. Clinical and biological annotations of the samples have been declared to the CNIL (Comité National Informatique et Libertés).

#### Patient-derived xenograft

Viably frozen PDX cells were thawed, washed in PBS/2% FBS before resuspension in Matrigel:PBS (1:2) and subcutaneous injection of 0.5×106 cells into the left flank of a NGS mouse (Janvier Labs, Le Genest-Saint-Isle, France). Mice were housed under pathogen-free conditions in an animal room at constant temperature (20°C-22°C), with a 12-hour light/dark cycle and free access to food and water. Mice body weights and tumor volumes were measured 3 times a week with calipers, using the following formula: $(\text{length} \times \text{width}^2 \times \pi)/6$. At the end of the experiment, mice (8 per group) were humanely sacrificed. All animal procedures were performed following INSERM principles and guidelines. The protocol was approved by Midi-Pyrénées Ethics Committee on Animal Experimentation and conducted in accordance with institutional guidelines and the 2010/63/EU Directive under protocol number DAP-APAFiS-2021100714596279-, CEEA122 agreement: 2021100714596279_v9.

#### Multiplex Immunofluorescence (mIF) BOND RX

The BOND RX staining platform (LEICA Biosystems, Nussloch, Germany) was used to automate the mIF staining procedure on 4µm FFPE tissue sections (software version 7.0.9.267 RX). After baking and dewaxing, tissue slides were heat-pretreated for 20 minutes at 100°C using ER2 pretreatment solution (pH9, LEICA Biosystems, Nussloch, Germany). The slides were blocked for endogenous peroxidase activity using the Discovery inhibitor (15 minutes at room temperature) (07017944001, Roche Diagnostics, Basel, Switzerland), and stained for 4-plex immunofluorescence using the OPAL Technology (AKOYA Biosciences, Marlborough, USA) with sequential denaturation by ER1 pretreatment buffer (pH6, LEICA Biosystems, Nussloch, Germany), for 20 minutes at 97°C between each antibody-staining cycle. Primary antibodies were incubated in the following sequence: ALK1 (clone ALK01, Ready-to-use, Roche Diagnostics, Basel, Switzerland), CD163 (clone MRQ-26, 05973929001, Ready-to-use, Roche Diagnostics, Basel, Switzerland) and CD68 (clone PGM-1, 1/100 in Envison Flex Diluent, Agilent Technologies, Santa Clara, CA, USA). Four OPAL dyes with excitation and emission wavelengths compatible with our whole slide imaging system were used: OPAL Polaris 480, OPAL 520, OPAL 570 (1/300 in 1x Plus Automation Amplification Diluent, AKOYA Biosciences, Marlborough, USA) and OPAL 690 (1/150 in 1x Plus Automation Amplification Diluent, AKOYA Biosciences, Marlborough, USA). The tissue slides were counterstained using Spectral DAPI (AKOYA Biosciences, Marlborough, USA) and mounted with Invitrogen ProLong Gold Antifade Mounting medium (Life Technologies, ThermoFisher Scientific, California, USA).

Multispectral fluorescence imaging was performed using an AxioScan Z1 (Carl Zeiss Microscopy, Oberkochen, Germany) whole-slide scanner with appropriate narrow band-pass excitation and emission filters and specific dichroic mirrors (Semrock Inc., Rochester, NY, USA), and with a multi-channel solid-state light engine (Colibri 7, Carl Zeiss Microscopy, Oberkochen, Germany) equipped with 7 LEDs covering the entire visible spectrum from UV to far-red (370 – 648 nm). Analog to digital image sampling was performed at 16-bit (65 536 grey levels, 37 000:1 dynamic range) with a high-resolution scientific complementary metal oxide semiconductor (sCMOS sensor with 2 048 x 2 048 cells of size 6.5 x 6.5 μm each)

Peltier-cooled monochrome camera (Orca Flash 4.0 V3, Hamamatsu Photonics K.K., Japan), to achieve a final scan resolution of 0.32 μm/pixel.

### Supplementary Data

**Supplementary Table 1.** Characteristics of patients with ALCL (n = 69 patients with 66 affymetrix and 39 RNAseq samples) stratified according to relapse status or morphological type.

| ALCL patients data | | | | | |
|---|---|---|---|---|---|
| **Microarray** | **RNAseq** | **Patients** | **Mophological type** | **Relapsing Group** | **ALCL subtype** |
| ✓ | | ALCL01 | CT | Relapse | ALCL ALK(+) |
| ✓ | | ALCL02 | CT | No Relapse | ALCL ALK(+) |
| ✓ | | ALCL03 | CT | Relapse | ALCL ALK(+) |
| ✓ | ✓ | **ALCL04** | nonCT | Relapse | ALCL ALK(+) |
| ✓ | ✓ | **ALCL05** | nonCT | Relapse | ALCL ALK(+) |
| ✓ | ✓ | **ALCL06** | CT | No Relapse | ALCL ALK(+) |
| ✓ | | ALCL07 | nonCT | No Relapse | ALCL ALK(+) |
| ✓ | ✓ | **ALCL08** | CT | No Relapse | ALCL ALK(+) |
| ✓ | ✓ | **ALCL09** | CT | No Relapse | ALCL ALK(+) |
| ✓ | | ALCL10 | nonCT | Relapse | ALCL ALK(+) |
| ✓ | ✓ | **ALCL11** | CT | Relapse | ALCL ALK(+) |
| ✓ | ✓ | **ALCL12** | CT | No Relapse | ALCL ALK(+) |
| ✓ | | ALCL13 | NA | Relapse | ALCL ALK(-) |
| ✓ | | ALCL18 | nonCT | No Relapse | ALCL ALK(+) |
| ✓ | ✓ | **ALCL19** | CT | Relapse | ALCL ALK(+) |
| ✓ | | ALCL21 | CT | Relapse | ALCL ALK(+) |
| ✓ | ✓ | **ALCL22** | nonCT | Relapse | ALCL ALK(+) |
| ✓ | | ALCL23 | CT | No Relapse | ALCL ALK(+) |
| ✓ | ✓ | **ALCL24** | CT | Relapse | ALCL ALK(+) |
| ✓ | ✓ | **ALCL25** | nonCT | Relapse | ALCL ALK(+) |
| ✓ | | ALCL26 | nonCT | Relapse | ALCL ALK(+) |
| ✓ | ✓ | **ALCL27** | nonCT | Relapse | ALCL ALK(+) |
| ✓ | | ALCL28 | nonCT | NA | ALCL ALK(+) |
| ✓ | ✓ | **ALCL29** | CT | No Relapse | ALCL ALK(+) |

| | | | | | |
|---|---|---|---|---|---|
| ✓ | ✓ | **ALCL30** | nonCT | No Relapse | ALCL ALK(+) |
| ✓ | ✓ | **ALCL31** | CT | Relapse | ALCL ALK(+) |
| ✓ | | ALCL32 | nonCT | NA | ALCL ALK(+) |
| ✓ | | ALCL33 | nonCT | NA | ALCL ALK(+) |
| ✓ | | ALCL34 | NA | Relapse | ALCL ALK(-) |
| ✓ | ✓ | **ALCL35** | CT | No Relapse | ALCL ALK(+) |
| ✓ | ✓ | **ALCL36** | CT | No Relapse | ALCL ALK(+) |
| ✓ | ✓ | **ALCL37** | CT | Relapse | ALCL ALK(+) |
| ✓ | ✓ | **ALCL38** | nonCT | Relapse | ALCL ALK(+) |
| ✓ | ✓ | **ALCL39** | nonCT | No Relapse | ALCL ALK(+) |
| ✓ | | ALCL40 | nonCT | Relapse | ALCL ALK(+) |
| ✓ | | ALCL41 | NA | No Relapse | ALCL ALK(-) |
| ✓ | ✓ | **ALCL42** | CT | Relapse | ALCL ALK(+) |
| ✓ | ✓ | **ALCL43** | CT | No Relapse | ALCL ALK(+) |
| ✓ | | ALCL45 | nonCT | Relapse | ALCL ALK(+) |
| ✓ | | ALCL48 | NA | No Relapse | ALCL ALK(-) |
| ✓ | | ALCL50 | nonCT | No Relapse | ALCL ALK(+) |
| ✓ | ✓ | **ALCL51** | nonCT | Relapse | ALCL ALK(+) |
| ✓ | ✓ | **ALCL52** | CT | Relapse | ALCL ALK(+) |
| ✓ | ✓ | **ALCL53** | nonCT | Relapse | ALCL ALK(+) |
| ✓ | ✓ | **ALCL55** | nonCT | Relapse | ALCL ALK(+) |
| ✓ | ✓ | **ALCL56** | CT | No Relapse | ALCL ALK(+) |
| ✓ | ✓ | **ALCL57** | nonCT | Relapse | ALCL ALK(+) |
| ✓ | ✓ | **ALCL58** | CT | No Relapse | ALCL ALK(+) |
| ✓ | ✓ | **ALCL59** | nonCT | Relapse | ALCL ALK(+) |
| ✓ | | ALCL60 | CT | Relapse | ALCL ALK(+) |
| ✓ | | ALCL61 | nonCT | No Relapse | ALCL ALK(+) |
| ✓ | ✓ | **ALCL62** | CT | No Relapse | ALCL ALK(+) |
| ✓ | ✓ | **ALCL63** | nonCT | No Relapse | ALCL ALK(+) |
| ✓ | | ALCL64 | nonCT | No Relapse | ALCL ALK(+) |

| | | | | | |
|---|---|---|---|---|---|
| ✓ | | ALCL65 | CT | No Relapse | ALCL ALK(+) |
| ✓ | | ALCL66 | nonCT | NA | ALCL ALK(+) |
| ✓ | | ALCL67 | nonCT | No Relapse | ALCL ALK(+) |
| ✓ | | ALCL68 | NA | Relapse | ALCL ALK(-) |
| ✓ | | ALCL69 | CT | No Relapse | ALCL ALK(+) |
| ✓ | | ALCL70 | nonCT | Relapse | ALCL ALK(+) |
| ✓ | ✓ | **ALCL72** | CT | No Relapse | ALCL ALK(+) |
| ✓ | ✓ | **ALCL73** | nonCT | No Relapse | ALCL ALK(+) |
| ✓ | ✓ | **ALCL74** | nonCT | No Relapse | ALCL ALK(+) |
| ✓ | | ALCL80 | NA | No Relapse | ALCL ALK(-) |
| ✓ | | ALCL81 | nonCT | Relapse | ALCL ALK(+) |
| ✓ | ✓ | **ALCL82 (P_NR_V01)** | CT | No Relapse | ALCL ALK(+) |
| | ✓ | **ALCL83 (P_NR_V02)** | CT | No Relapse | ALCL ALK(+) |
| | ✓ | **ALCL84 (P_NR_V08)** | NA | No Relapse | ALCL ALK(+) |
| | ✓ | **ALCL89 (P_NR_V16)** | CT | No Relapse | ALCL ALK(+) |

**Abbreviations :** ALCL = Anaplastic Large Cell Lymphoma; CT = Common-Type variant; nonCT = non-Common-Type variant.

**Supplementary Table 2.** List of macrophage gene signatures.

| name | # genes | description | collections | source organism | contributor | common_list* |
|---|---|---|---|---|---|---|
| **WP_MACROPHAGE_MARKERS** | **9** | **Macrophage markers** | **C2 CP** | **Homo sapiens** | **WikiPathways** | **TRUE** |
| **GOBP_MACROPHAGE_ACTIVATION** | **110** | **A change in morphology and behavior of a macrophage resulting from exposure to a cytokine, chemokine, cellular ligand, or soluble factor. [GOC:mgi_curators, ISBN:0781735149, PMID:14506301]** | **C5 GO** | **Homo sapiens** | **Gene Ontology Consortium** | **TRUE** |
| **GOBP_MACROPHAGE_ACTIVATION_INVOLVED_IN_IMMUNE_RESPONSE** | **19** | **A change in morphology and behavior of a macrophage resulting from exposure to a cytokine, chemokine, cellular ligand, or soluble factor, leading to the initiation or perpetuation of an immune response. [GOC:add, ISBN:0781735149]** | **C5 GO** | **Homo sapiens** | **Gene Ontology Consortium** | **TRUE** |
| GOBP_REGULATION_OF_MACROPHAGE_APOPTOTIC_PROCESS | **11** | Any process that modulates the frequency, rate or extent of macrophage apoptotic process. [GOC:BHF, GOC:mtg_apoptosis] | C5 GO | Homo sapiens | Gene Ontology Consortium | FALSE |
| GOBP_MACROPHAGE_CHEMOTAXIS | **41** | The movement of a macrophage in response to an external stimulus. [GOC:jid] | C5 GO | Homo sapiens | Gene Ontology Consortium | FALSE |
| GOBP_MACROPHAGE_COLONY_STIMULATING_FACTOR_PRODUCTION | **7** | The appearance of macrophage colony-stimulating factor due to biosynthesis or secretion following a cellular stimulus, resulting in an increase in its intracellular or extracellular levels. [GOC:BHF, GOC:vk] | C5 GO | Homo sapiens | Gene Ontology Consortium | FALSE |
| GOBP_MACROPHAGE_COLONY_STIMULATING_FACTOR_SIGNALING_PATHWAY | **7** | The series of molecular signals initiated by the binding of the cytokine macrophage colony-stimulating factor (M-CSF) to its receptor on the surface of a target cell, and ending with the regulation of a downstream cellular process, e.g. transcription. [GOC:signaling, GOC:uh, PMID:12138890, Wikipedia:Macrophage_colony-stimulating_factor] | C5 GO | Homo sapiens | Gene Ontology Consortium | FALSE |

| | | | | | | |
|---|---|---|---|---|---|---|
| GOBP_MACROPHAGE_CYTOKINE_PRODUCTION | **37** | The appearance of a macrophage cytokine due to biosynthesis or secretion following a cellular stimulus, resulting in an increase in its intracellular or extracellular levels. [GOC:BHF, GOC:dph, GOC:rl, GOC:tb] | C5 GO | Homo sapiens | Gene Ontology Consortium | FALSE |
| GOBP_REGULATION_OF_MACROPHAGE_DIFFERENTIATION | **25** | Any process that modulates the frequency, rate or extent of macrophage differentiation. [GOC:go_curators] | C5 GO | Homo sapiens | Gene Ontology Consortium | FALSE |
| GOBP_MACROPHAGE_FUSION | **6** | The binding and fusion of a macrophage to one or more other cells to form a multinucleated cell. [GOC:sl] | C5 GO | Homo sapiens | Gene Ontology Consortium | FALSE |
| GOBP_MACROPHAGE_INFLAMMATORY_PROTEIN_1_ALPHA_PRODUCTION | **6** | The appearance of macrophage inflammatory protein 1 alpha due to biosynthesis or secretion following a cellular stimulus, resulting in an increase in its intracellular or extracellular levels. [GOC:add, GOC:rv] | C5 GO | Homo sapiens | Gene Ontology Consortium | FALSE |
| GOBP_MACROPHAGE_MIGRATION | **61** | The orderly movement of a macrophage from one site to another. [GO_REF:0000091, GOC:TermGenie, PMID:25749876] | C5 GO | Homo sapiens | Gene Ontology Consortium | FALSE |
| **GOBP_MACROPHAGE_PROLIFERATION** | **13** | **The expansion of a macrophage population by cell division. [GOC:dph, PMID:12614284, PMID:19466391]** | **C5 GO** | **Homo sapiens** | **Gene Ontology Consortium** | **TRUE** |
| GOBP_POSITIVE_REGULATION_OF_MACROPHAGE_ACTIVATION | **30** | Any process that stimulates, induces or increases the rate of macrophage activation. [GOC:jl] | C5 GO | Homo sapiens | Gene Ontology Consortium | FALSE |
| GOBP_POSITIVE_REGULATION_OF_MACROPHAGE_CHEMOTAXIS | **19** | Any process that increases the rate, frequency or extent of macrophage chemotaxis. Macrophage chemotaxis is the movement of a macrophage in response to an external stimulus. [GOC:BHF, GOC:dph, GOC:tb] | C5 GO | Homo sapiens | Gene Ontology Consortium | FALSE |
| GOBP_POSITIVE_REGULATION_OF_MACROPHAGE_CYTOKINE_PRODUCTION | **24** | Any process that increases the rate, frequency or extent of macrophage cytokine production. Macrophage cytokine production is the appearance of a chemokine due to biosynthesis or secretion following a cellular stimulus, resulting in an increase in its intracellular or | C5 GO | Homo sapiens | Gene Ontology Consortium | FALSE |

| | | | | | | |
|---|---|---|---|---|---|---|
| | | extracellular levels. [GOC:dph, GOC:tb] | | | | |
| GOBP_POSITIVE_REGULATION_OF_MACROPHAGE_DIFFERENTIATION | **16** | Any process that activates or increases the frequency, rate or extent of macrophage differentiation. [GOC:go_curators] | C5 GO | Homo sapiens | Gene Ontology Consortium | FALSE |
| GOBP_POSITIVE_REGULATION_OF_MACROPHAGE_MIGRATION | **26** | Any process that activates or increases the frequency, rate or extent of macrophage migration. [GO_REF:0000058, GOC:TermGenie, PMID:25749876] | C5 GO | Homo sapiens | Gene Ontology Consortium | FALSE |
| GOBP_REGULATION_OF_MACROPHAGE_ACTIVATION | **62** | Any process that modulates the frequency or rate of macrophage activation. [GOC:jl] | C5 GO | Homo sapiens | Gene Ontology Consortium | FALSE |
| GOBP_REGULATION_OF_MACROPHAGE_CHEMOTAXIS | **28** | Any process that modulates the rate, frequency or extent of macrophage chemotaxis. Macrophage chemotaxis is the movement of a macrophage in response to an external stimulus. [GOC:BHF, GOC:dph, GOC:tb] | C5 GO | Homo sapiens | Gene Ontology Consortium | FALSE |
| GOBP_REGULATION_OF_MACROPHAGE_FUSION | **5** | Any process that modulates the frequency, rate or extent of macrophage fusion. [GOC:mah] | C5 GO | Homo sapiens | Gene Ontology Consortium | FALSE |
| GOBP_REGULATION_OF_MACROPHAGE_MIGRATION | **45** | Any process that modulates the frequency, rate or extent of macrophage migration. [GO_REF:0000058, GOC:TermGenie, PMID:25749876] | C5 GO | Homo sapiens | Gene Ontology Consortium | FALSE |
| GOBP_REGULATION_OF_MACROPHAGE_PROLIFERATION | **7** | Any process that modulates the frequency, rate or extent of macrophage proliferation. [GOC:BHF, GOC:BHF_miRNA, GOC:rph] | C5 GO | Homo sapiens | Gene Ontology Consortium | FALSE |
| **GOBP_MACROPHAGE_APOPTOTIC_PROCESS** | **11** | | | **Homo sapiens** | | **TRUE** |
| **GOBP_MACROPHAGE_DIFFERENTIATION** | **47** | | | **Homo sapiens** | | **TRUE** |

* TRUE = signature score significatively different between Common Type and non-Common Type variants from both Affymetrix and RNAseq data.

**Supplementary Table 3.** List of macrophage gene signatures.

| name | # genes | description | collections | source organism | contributor | common_list * |
|---|---|---|---|---|---|---|
| **GOBP_MONOCYTE_ACTIVATION** | **12** | **The change in morphology and behavior of a monocyte resulting from exposure to a cytokine, chemokine, cellular ligand, or soluble factor. [GOC:mgi_curators, ISBN:0781735149]** | **C5 GO** | **Homo sapiens** | **Gene Ontology Consortium** | **TRUE** |
| GOBP_MONOCYTE_AGGREGATION | 5 | The adhesion of one monocyte to one or more other monocytes via adhesion molecules. [GOC:sl, PMID:12972508] | C5 GO | Homo sapiens | Gene Ontology Consortium | FALSE |
| **GOBP_MONOCYTE_CHEMOTACTIC_PROTEIN_1_PRODUCTION** | **21** | **The appearance of monocyte chemotactic protein-1 due to biosynthesis or secretion following a cellular stimulus, resulting in an increase in its intracellular or extracellular levels. [GOC:add, GOC:rv]** | **C5 GO** | **Homo sapiens** | **Gene Ontology Consortium** | **TRUE** |
| **GOBP_MONOCYTE_CHEMOTAXIS** | **71** | **The movement of a monocyte in response to an external stimulus. [GOC:add, PMID:11696603, PMID:15173832]** | **C5 GO** | **Homo sapiens** | **Gene Ontology Consortium** | **TRUE** |
| **GOBP_MONOCYTE_EXTRAVASATION** | **10** | **The migration of a monocyte from the blood vessels into the surrounding tissue. [CL:0000576, GOC:BHF, PMID:10657654]** | **C5 GO** | **Homo sapiens** | **Gene Ontology Consortium** | **TRUE** |
| **GOBP_REGULATION_OF_MONOCYTE_CHEMOTAXIS** | **29** | **Any process that modulates the frequency, rate, or extent of monocyte chemotaxis. [GOC:dph, GOC:tb]** | **C5 GO** | **Homo sapiens** | **Gene Ontology Consortium** | **TRUE** |
| GOBP_REGULATION_OF_MONOCYTE_DIFFERENTIATION | 20 | Any process that modulates the frequency, rate or extent of monocyte differentiation. [GOC:go_curators] | C5 GO | Homo sapiens | Gene Ontology Consortium | FALSE |
| **BIOCARTA_MONOCYTE_PATHWAY** | **11** | **Monocyte and its Surface Molecules** | **C2 CP** | **Homo sapiens** | **BioCarta** | **TRUE** |
| GOBP_MONOCYTE_ACTIVATION | 12 | The change in morphology and behavior of a monocyte resulting from exposure to a cytokine, chemokine, cellular ligand, or soluble factor. [GOC:mgi_curators, ISBN:0781735149] | C5 GO | Homo sapiens | Gene Ontology Consortium | FALSE |

* TRUE = signature score significatively different between Common Type and non-Common Type variants from both Affymetrix and RNAseq data

**Supplementary Table 4.** Table of macrophage and monocyte genes differentially expressed in common type morphology patients versus non-common type ones, for both Affymetrix (left) and RNAseq (right) datasets. Genes underlined in grey are from monocyte gene list and in blue are from macrophage gene list.

| **Affymetrix differential expression analysis** | | | | | | **RNAseq differential expression analysis** | | | | |
|---|---|---|---|---|---|---|---|---|---|---|
| **probe** | **gene** | **logFC** | **P.Value** | **adj.P.Val** | | **target_id** | **ext_gene** | **pval** | **qval** | **b** |
| 225353_s_at | C1QC | -1.18 | 8.83E-06 | 1.41E-04 | | **ENSG00000275385.2** | **CCL18** | **7.29E-04** | **6.41E-03** | **-1.84** |
| **32128_at** | **CCL18** | **1.76** | **6.40E-04** | **4.40E-03** | | **ENSG00000172724.12** | **CCL19** | **4.20E-04** | **4.25E-03** | **2.50** |
| **209924_at** | **CCL18** | **1.72** | **4.34E-04** | **3.23E-03** | | ENSG00000136999.5 | **CCN3** | 1.25E-04 | 1.73E-03 | -1.42 |
| **210072_at** | **CCL19** | **-3.00** | **4.79E-06** | **8.62E-05** | | **ENSG00000172243.18** | **CLEC7A** | **1.22E-05** | **3.12E-04** | **1.07** |
| 207794_at | CCR2 | -1.26 | 2.91E-05 | 3.66E-04 | | ENSG00000164400.6 | **CSF2** | 2.51E-04 | 2.91E-03 | 1.37 |
| 206978_at | CCR2 | -1.52 | 2.31E-06 | 4.84E-05 | | **ENSG00000169245.6** | **CXCL10** | **7.16E-03** | **3.55E-02** | **1.25** |
| 209619_at | CD74 | -1.12 | 1.28E-09 | 1.88E-07 | | **ENSG00000111537.5** | **IFNG** | **2.26E-05** | **4.93E-04** | **1.61** |
| 1567628_at | CD74 | -2.07 | 7.71E-12 | 8.04E-09 | | ENSG00000157368.12 | **IL34** | 4.27E-04 | 4.30E-03 | 1.30 |
| 204039_at | CEBPA | -1.24 | 6.04E-08 | 2.88E-06 | | ENSG00000160593.19 | **JAML** | 8.66E-09 | 1.81E-06 | 1.34 |
| **1554406_a_at** | **CLEC7A** | **-1.44** | **2.89E-06** | **5.74E-05** | | **ENSG00000129988.6** | **LBP** | **7.72E-03** | **3.75E-02** | **-1.27** |
| **221698_s_at** | **CLEC7A** | **-1.55** | **7.87E-08** | **3.56E-06** | | **ENSG00000090382.7** | **LYZ** | **1.26E-04** | **1.73E-03** | **1.30** |

| | | | | |
|---|---|---|---|---|
| **1555214_a_at** | **CLEC7A** | **-1.57** | **4.29E-06** | **7.91E-05** |
| **1555756_a_at** | **CLEC7A** | **-1.87** | **6.07E-07** | **1.70E-05** |
| **204533_at** | **CXCL10** | **-2.08** | **3.89E-05** | **4.64E-04** |
| 228410_at | GAB3 | -1.24 | 1.16E-06 | 2.84E-05 |
| 218469_at | GREM1 | -1.13 | 1.29E-02 | 4.88E-02 |
| 221491_x_at | HLA-DRB1 /// (…) | -1.51 | 5.11E-04 | 3.69E-03 |
| 215193_x_at | HLA-DRB1 /// (…) | -1.16 | 7.92E-09 | 6.27E-07 |
| 209312_x_at | HLA-DRB1 /// (…) | -1.10 | 2.08E-09 | 2.59E-07 |
| 204670_x_at | HLA-DRB1 /// (…) | -1.12 | 3.61E-09 | 3.77E-07 |
| **210354_at** | **IFNG** | **-2.10** | **4.37E-05** | **5.10E-04** |
| 205992_s_at | IL15 | -1.04 | 2.01E-06 | 4.34E-05 |
| 213475_s_at | ITGAL | -1.32 | 1.23E-07 | 5.06E-06 |
| 1554240_a_at | ITGAL | -1.59 | 2.14E-07 | 7.73E-06 |

| | | | | |
|---|---|---|---|---|
| ENSG00000207975.1 | **MIR181B1** | 1.79E-05 | 4.20E-04 | 1.18 |
| ENSG00000125144.14 | **MT1G** | 3.98E-05 | 7.50E-04 | -1.35 |
| ENSG00000130584.12 | **ZBTB46** | 2.91E-07 | 2.14E-05 | -1.29 |

| | | | | |
|---|---|---|---|---|
| 1555349_a_at | ITGB2 | -1.08 | 3.43E-06 | 6.59E-05 |
| **214461_at** | **LBP** | **1.27** | **7.44E-03** | **3.18E-02** |
| **213975_s_at** | **LYZ** | **-1.33** | **1.07E-07** | **4.52E-06** |
| **1555745_a_at** | **LYZ** | **-2.43** | **3.96E-07** | **1.23E-05** |
| 204563_at | SELL | -1.34 | 3.64E-05 | 4.40E-04 |
| 219385_at | SLAMF8 | -1.28 | 7.23E-07 | 1.96E-05 |
| 219386_s_at | SLAMF8 | -1.40 | 9.84E-07 | 2.49E-05 |
| 223939_at | SUCNR1 | -1.04 | 2.94E-04 | 2.36E-03 |
| 204122_at | TYROBP | -1.37 | 5.73E-07 | 1.63E-05 |

**Supplementary Table 5.** Transcriptomic markers from MCP-Counter reduced list (selection on cell types *Neutrophils* and *Monocytes*).

| **Gene** | **Population** | **reduce_signature** |
|---|---|---|
| CA4 | Neutrophils | Yes |
| CEACAM3 | Neutrophils | Yes |
| CXCR1 | Neutrophils | Yes |
| CXCR2 | Neutrophils | Yes |
| CYP4F3 | Neutrophils | Yes |
| FCGR3B | Neutrophils | Yes |
| HAL | Neutrophils | Yes |
| KCNJ15 | Neutrophils | Yes |
| LOC254896 /// TNFRSF10C | Neutrophils | Yes |
| MEGF9 | Neutrophils | Yes |
| SLC25A37 | Neutrophils | Yes |
| STEAP4 | Neutrophils | Yes |
| TECPR2 | Neutrophils | Yes |
| TLE3 | Neutrophils | Yes |
| TNFRSF10C | Neutrophils | Yes |
| VNN3 | Neutrophils | Yes |
| CSF1R | Monocytic lineage | Yes |
| KYNU | Monocytic lineage | Yes |
| PLA2G7 | Monocytic lineage | Yes |
| ADAP2 | Monocytic lineage | Yes |
| RASSF4 | Monocytic lineage | Yes |
| FPR3 | Monocytic lineage | Yes |
| TFEC | Monocytic lineage | Yes |

**Supplementary Table 6.** Significative macrophage signature genes from Affymetrix and RNAseq data analysis.

| gene_name | RNAseq_sign* | Affy_sign* | RNAseq&Affy_sign* |
|---|---|---|---|
| ADIPOQ | FALSE | FALSE | FALSE |
| APP | FALSE | FALSE | FALSE |
| BMP4 | FALSE | FALSE | FALSE |
| C1QC | FALSE | TRUE | FALSE |
| CASP8 | FALSE | FALSE | FALSE |
| CD14 | FALSE | FALSE | FALSE |
| CD163 | FALSE | FALSE | FALSE |
| CD4 | FALSE | FALSE | FALSE |
| CD68 | FALSE | FALSE | FALSE |
| CD74 | FALSE | TRUE | FALSE |
| CD83 | FALSE | FALSE | FALSE |
| CD86 | FALSE | FALSE | FALSE |
| CDC42 | FALSE | FALSE | FALSE |
| CDKN2A | FALSE | FALSE | FALSE |
| CEBPA | FALSE | TRUE | FALSE |
| CEBPE | FALSE | FALSE | FALSE |
| CLU | FALSE | FALSE | FALSE |
| CSF1 | FALSE | FALSE | FALSE |
| CSF1R | FALSE | FALSE | FALSE |
| **CSF2** | TRUE | FALSE | FALSE |
| CTSL | FALSE | FALSE | FALSE |
| CX3CL1 | FALSE | FALSE | FALSE |
| DYSF | FALSE | FALSE | FALSE |
| EIF2AK1 | FALSE | FALSE | FALSE |
| F3 | FALSE | FALSE | FALSE |
| FADD | FALSE | FALSE | FALSE |
| FER1L5 | FALSE | FALSE | FALSE |
| GAB3 | FALSE | TRUE | FALSE |
| GATA2 | FALSE | FALSE | FALSE |
| GBA | FALSE | FALSE | FALSE |
| GHSR | FALSE | FALSE | FALSE |
| GRN | FALSE | FALSE | FALSE |
| HAVCR2 | FALSE | FALSE | FALSE |

| | | | |
|---|---|---|---|
| HCLS1 | FALSE | FALSE | FALSE |
| HLA-DRB1 | FALSE | TRUE | FALSE |
| HMGB1 | FALSE | FALSE | FALSE |
| IFI35 | FALSE | FALSE | FALSE |
| **IFNG** | **TRUE** | **TRUE** | **TRUE** |
| IL15 | FALSE | TRUE | FALSE |
| IL31RA | FALSE | FALSE | FALSE |
| IL33 | FALSE | FALSE | FALSE |
| **IL34** | TRUE | FALSE | FALSE |
| INHA | FALSE | FALSE | FALSE |
| INHBA | FALSE | FALSE | FALSE |
| IRF3 | FALSE | FALSE | FALSE |
| IRF7 | FALSE | FALSE | FALSE |
| L3MBTL3 | FALSE | FALSE | FALSE |
| **LBP** | **TRUE** | **TRUE** | **TRUE** |
| LIF | FALSE | FALSE | FALSE |
| **LYZ** | **TRUE** | **TRUE** | **TRUE** |
| MAPK1 | FALSE | FALSE | FALSE |
| MAPK3 | FALSE | FALSE | FALSE |
| MEF2C | FALSE | FALSE | FALSE |
| MIR145 | FALSE | FALSE | FALSE |
| **MIR181B1** | TRUE | FALSE | FALSE |
| MIR223 | FALSE | FALSE | FALSE |
| MMP9 | FALSE | FALSE | FALSE |
| NKX2-3 | FALSE | FALSE | FALSE |
| NMI | FALSE | FALSE | FALSE |
| NOD2 | FALSE | FALSE | FALSE |
| NRROS | FALSE | FALSE | FALSE |
| PARP1 | FALSE | FALSE | FALSE |
| PF4 | FALSE | FALSE | FALSE |
| PLEKHO2 | FALSE | FALSE | FALSE |
| PRKCA | FALSE | FALSE | FALSE |
| PRKCE | FALSE | FALSE | FALSE |
| PTK2 | FALSE | FALSE | FALSE |
| PTPN2 | FALSE | FALSE | FALSE |
| RAC2 | FALSE | FALSE | FALSE |
| RB1 | FALSE | FALSE | FALSE |

| | | | |
|---|---|---|---|
| RIPK1 | FALSE | FALSE | FALSE |
| ROR2 | FALSE | FALSE | FALSE |
| SBNO2 | FALSE | FALSE | FALSE |
| SELENOS | FALSE | FALSE | FALSE |
| SIRT1 | FALSE | FALSE | FALSE |
| SPI1 | FALSE | FALSE | FALSE |
| SUCNR1 | FALSE | TRUE | FALSE |
| SYK | FALSE | FALSE | FALSE |
| TCP1 | FALSE | FALSE | FALSE |
| TGFB1 | FALSE | FALSE | FALSE |
| TICAM1 | FALSE | FALSE | FALSE |
| TLR2 | FALSE | FALSE | FALSE |
| TREM2 | FALSE | FALSE | FALSE |
| TREX1 | FALSE | FALSE | FALSE |
| TRIB1 | FALSE | FALSE | FALSE |
| TSPAN2 | FALSE | FALSE | FALSE |
| TYROBP | FALSE | TRUE | FALSE |
| VEGFA | FALSE | FALSE | FALSE |
| **ZBTB46** | TRUE | FALSE | FALSE |

*** FALSE = not present in the list of DE genes ; TRUE = present in the list of DE genes.**

**Abbreviations** : DE = differentially expressed.

**Supplementary Table 7.** Significative monocyte signature genes from Affymetrix and RNAseq data analysis.

| gene_name | RNAseq_sign* | Affy_sign* | RNAseq&Affy_sign* |
|---|---|---|---|
| ADAM10 | FALSE | FALSE | FALSE |
| ADAM9 | FALSE | FALSE | FALSE |
| ADIPOQ | FALSE | FALSE | FALSE |
| AGER | FALSE | FALSE | FALSE |
| AIF1 | FALSE | FALSE | FALSE |
| ANO6 | FALSE | FALSE | FALSE |
| ANXA1 | FALSE | FALSE | FALSE |
| APOD | FALSE | FALSE | FALSE |
| AZU1 | FALSE | FALSE | FALSE |
| C1QTNF3 | FALSE | FALSE | FALSE |
| CALCA | FALSE | FALSE | FALSE |
| CCL1 | FALSE | FALSE | FALSE |
| CCL11 | FALSE | FALSE | FALSE |
| CCL13 | FALSE | FALSE | FALSE |
| CCL14 | FALSE | FALSE | FALSE |
| CCL15 | FALSE | FALSE | FALSE |
| CCL16 | FALSE | FALSE | FALSE |
| CCL17 | FALSE | FALSE | FALSE |
| **CCL18** | **TRUE** | **TRUE** | **TRUE** |
| **CCL19** | **TRUE** | **TRUE** | **TRUE** |
| CCL2 | FALSE | FALSE | FALSE |
| CCL20 | FALSE | FALSE | FALSE |
| CCL21 | FALSE | FALSE | FALSE |
| CCL22 | FALSE | FALSE | FALSE |
| CCL23 | FALSE | FALSE | FALSE |
| CCL24 | FALSE | FALSE | FALSE |
| CCL25 | FALSE | FALSE | FALSE |

| | | | |
|---|---|---|---|
| CCL26 | FALSE | FALSE | FALSE |
| CCL3 | FALSE | FALSE | FALSE |
| CCL3L1 | FALSE | FALSE | FALSE |
| CCL3L3 | FALSE | FALSE | FALSE |
| CCL4 | FALSE | FALSE | FALSE |
| CCL5 | FALSE | FALSE | FALSE |
| CCL7 | FALSE | FALSE | FALSE |
| CCL8 | FALSE | FALSE | FALSE |
| **CCN3** | **TRUE** | FALSE | FALSE |
| CCR1 | FALSE | FALSE | FALSE |
| CCR2 | FALSE | TRUE | FALSE |
| CD33 | FALSE | FALSE | FALSE |
| CD44 | FALSE | FALSE | FALSE |
| CD47 | FALSE | FALSE | FALSE |
| **CLEC7A** | **TRUE** | **TRUE** | **TRUE** |
| CREB3 | FALSE | FALSE | FALSE |
| CSF1 | FALSE | FALSE | FALSE |
| CX3CL1 | FALSE | FALSE | FALSE |
| **CXCL10** | **TRUE** | **TRUE** | **TRUE** |
| CXCL12 | FALSE | FALSE | FALSE |
| CXCL17 | FALSE | FALSE | FALSE |
| DEFB104A | FALSE | FALSE | FALSE |
| DEFB104B | FALSE | FALSE | FALSE |
| DEFB124 | FALSE | FALSE | FALSE |
| DUSP1 | FALSE | FALSE | FALSE |
| DYSF | FALSE | FALSE | FALSE |
| ERBIN | FALSE | FALSE | FALSE |
| FER1L5 | FALSE | FALSE | FALSE |
| FLT1 | FALSE | FALSE | FALSE |
| FOXP1 | FALSE | FALSE | FALSE |

| | | | |
|---|---|---|---|
| FPR2 | FALSE | FALSE | FALSE |
| GREM1 | FALSE | TRUE | FALSE |
| GSTP1 | FALSE | FALSE | FALSE |
| HMGB1 | FALSE | FALSE | FALSE |
| HYAL2 | FALSE | FALSE | FALSE |
| ICAM1 | FALSE | FALSE | FALSE |
| IL1B | FALSE | FALSE | FALSE |
| IL6 | FALSE | FALSE | FALSE |
| IL6R | FALSE | FALSE | FALSE |
| ITGA4 | FALSE | FALSE | FALSE |
| ITGAL | FALSE | TRUE | FALSE |
| ITGAM | FALSE | FALSE | FALSE |
| ITGB1 | FALSE | FALSE | FALSE |
| ITGB2 | FALSE | TRUE | FALSE |
| **JAML** | **TRUE** | **FALSE** | **FALSE** |
| LGALS3 | FALSE | FALSE | FALSE |
| LGALS9 | FALSE | FALSE | FALSE |
| LGMN | FALSE | FALSE | FALSE |
| LILRB4 | FALSE | FALSE | FALSE |
| LYN | FALSE | FALSE | FALSE |
| MCOLN2 | FALSE | FALSE | FALSE |
| MIR92A1 | FALSE | FALSE | FALSE |
| MOSPD2 | FALSE | FALSE | FALSE |
| MSMP | FALSE | FALSE | FALSE |
| **MT1G** | **TRUE** | **FALSE** | **FALSE** |
| NBL1 | FALSE | FALSE | FALSE |
| NOD2 | FALSE | FALSE | FALSE |
| NR1H4 | FALSE | FALSE | FALSE |
| PDGFB | FALSE | FALSE | FALSE |
| PDGFD | FALSE | FALSE | FALSE |

| | | | |
|---|---|---|---|
| PECAM1 | FALSE | FALSE | FALSE |
| PLA2G7 | FALSE | FALSE | FALSE |
| PLCB1 | FALSE | FALSE | FALSE |
| PTPRO | FALSE | FALSE | FALSE |
| RPS19 | FALSE | FALSE | FALSE |
| S100A12 | FALSE | FALSE | FALSE |
| S100A14 | FALSE | FALSE | FALSE |
| S100A7 | FALSE | FALSE | FALSE |
| SELE | FALSE | FALSE | FALSE |
| SELENOK | FALSE | FALSE | FALSE |
| SELL | FALSE | TRUE | FALSE |
| SELP | FALSE | FALSE | FALSE |
| SERPINE1 | FALSE | FALSE | FALSE |
| SIRPA | FALSE | FALSE | FALSE |
| SLAMF8 | FALSE | TRUE | FALSE |
| SLIT2 | FALSE | FALSE | FALSE |
| SOCS5 | FALSE | FALSE | FALSE |
| SPACA3 | FALSE | FALSE | FALSE |
| SYK | FALSE | FALSE | FALSE |
| TNFRSF11A | FALSE | FALSE | FALSE |
| TNFSF11 | FALSE | FALSE | FALSE |
| TNFSF18 | FALSE | FALSE | FALSE |
| TWIST1 | FALSE | FALSE | FALSE |
| XCL1 | FALSE | FALSE | FALSE |
| XCL2 | FALSE | FALSE | FALSE |

*** FALSE = not present in the list of DE genes ; TRUE = present in the list of DE genes.**

**Abbreviations** : DE = differentially expressed

**Supplementary Figure S1: microenvironment cell signatures of ALCL versus AITL. (A)** The heatmap represents the Z-score (normalized cell-type score) of the 22 different immune cell types from CibersortX and 2 additional stromal cell types from MCP-Counter as in Figure 1. 161 samples are included : 78 ALCL (ALK+ and ALK-) and 83 AITL. The red frame highlights the cell subtypes enriched in ALCL and the green arrows highlight the AITL predominant subtypes. **(B)** The boxplot shows the different top enriched cell types into ALCL (grey frames) or AITL (green frames). Fibroblasts and Endothelial cells score scales are defined by MCP-Counter method while the others depend on the estimated proportions by CIBERSORTx. The p-values are calculated with a Wilcoxon test corrected by the Hochberg method. We kept p-val < 1e10-6 for graphical representation.

PTCL = Peripheral T-Cell Lymphoma; ALCL = Anaplastic Large Cell Lymphoma; AITL = Angioimmunoblastic T-Cell Lymphoma. In boxplots the center line represents the median, boxes indicate the interquartile range, the whiskers show the 1.5 interquartile range.

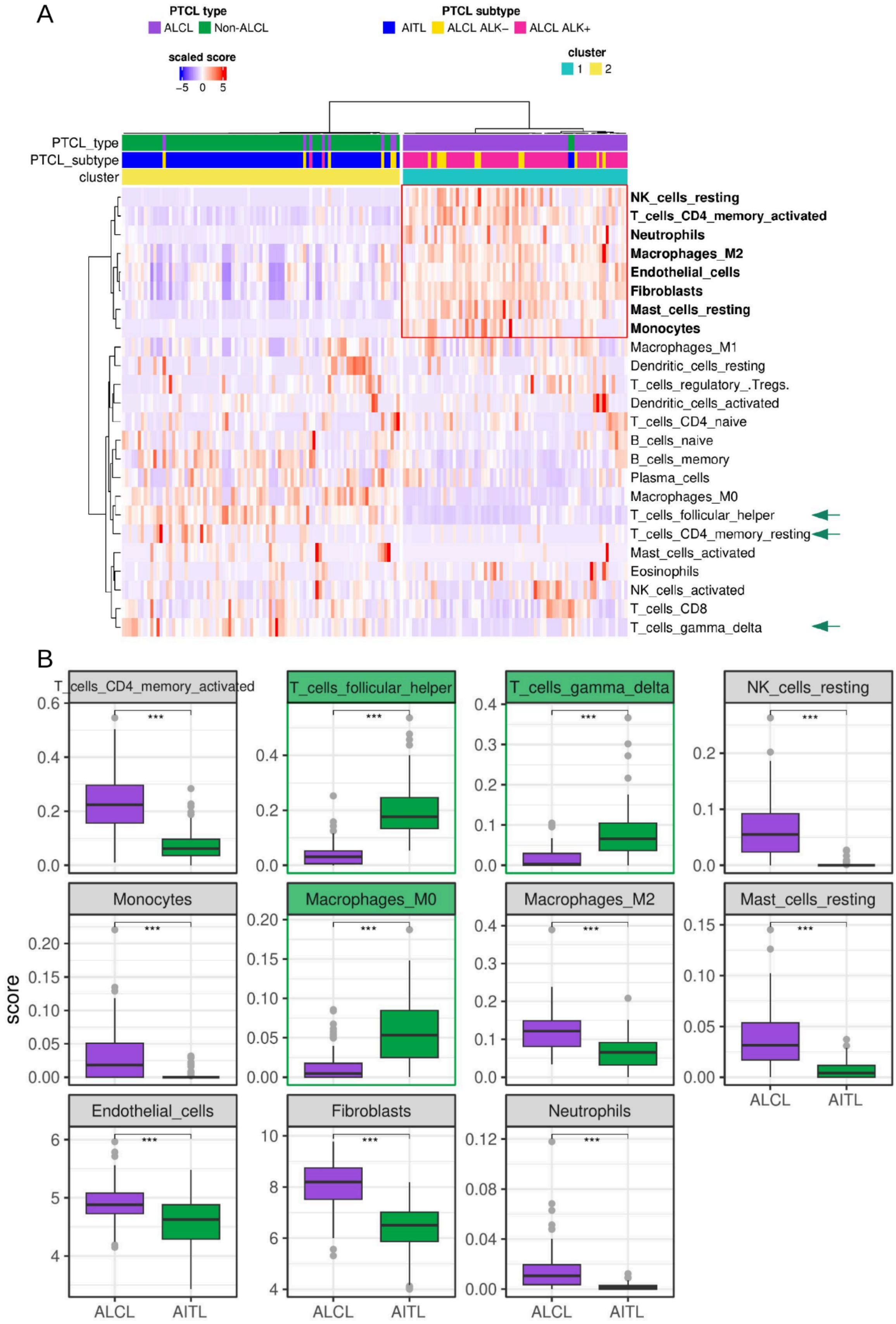
A
PTCL type
ALCL
Non-ALCL
PTCL subtype
AITL
ALCL ALK−
ALCL ALK+
scaled score
−5
0
5
cluster
1
2
PTCL_type
PTCL_subtype
cluster
NK_cells_resting
T_cells_CD4_memory_activated
Neutrophils
Macrophages_M2
Endothelial_cells
Fibroblasts
Mast_cells_resting
Monocytes
Macrophages_M1
Dendritic_cells_resting
T_cells_regulatory_.Tregs.
Dendritic_cells_activated
T_cells_CD4_naive
B_cells_naive
B_cells_memory
Plasma_cells
Macrophages_M0
T_cells_follicular_helper
T_cells_CD4_memory_resting
Mast_cells_activated
Eosinophils
NK_cells_activated
T_cells_CD8
T_cells_gamma_delta
B
score
T_cells_CD4_memory_activated
T_cells_follicular_helper
T_cells_gamma_delta
NK_cells_resting
Monocytes
Macrophages_M0
Macrophages_M2
Mast_cells_resting
Endothelial_cells
Fibroblasts
Neutrophils
***
ALCL
AITL

**Supplementary Figure S2: Dendritic cells activated scores into ALCL patients morphology groups.** Dendritic cells activated are enriched into CT patients compared to nonCT ones. CT and nonCT variants groups include 28 and 32 patients respectively. The p-value was calculated with a Wilcoxon test corrected by the Hochberg method. CT = Common-Type variant; nonCT = non-Common-Type variant. In boxplots the center line represents the median, boxes indicate the interquartile range, the whiskers show the 1.5 interquartile range.

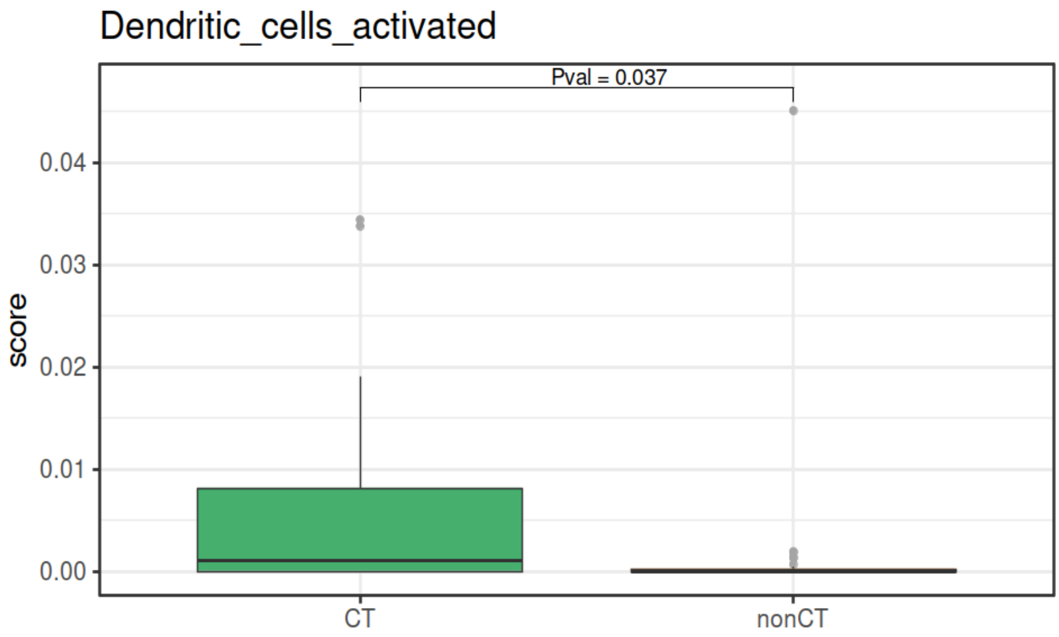

**Supplementary Figure S3: summary figure of ALCL patient samples included in the study (A) and the identified prognosis markers (B and C).** **(A)** ALCL patient Affymetrix samples at diagnosis (left) and RNA-seq samples (right) included in the study. **(B)** Numbers of macrophage signatures (blue) significatively different between CT and nonCT clusters and their number of associated differentially expressed genes (red). **(C)** Same as B for monocytes instead of Macrophages.

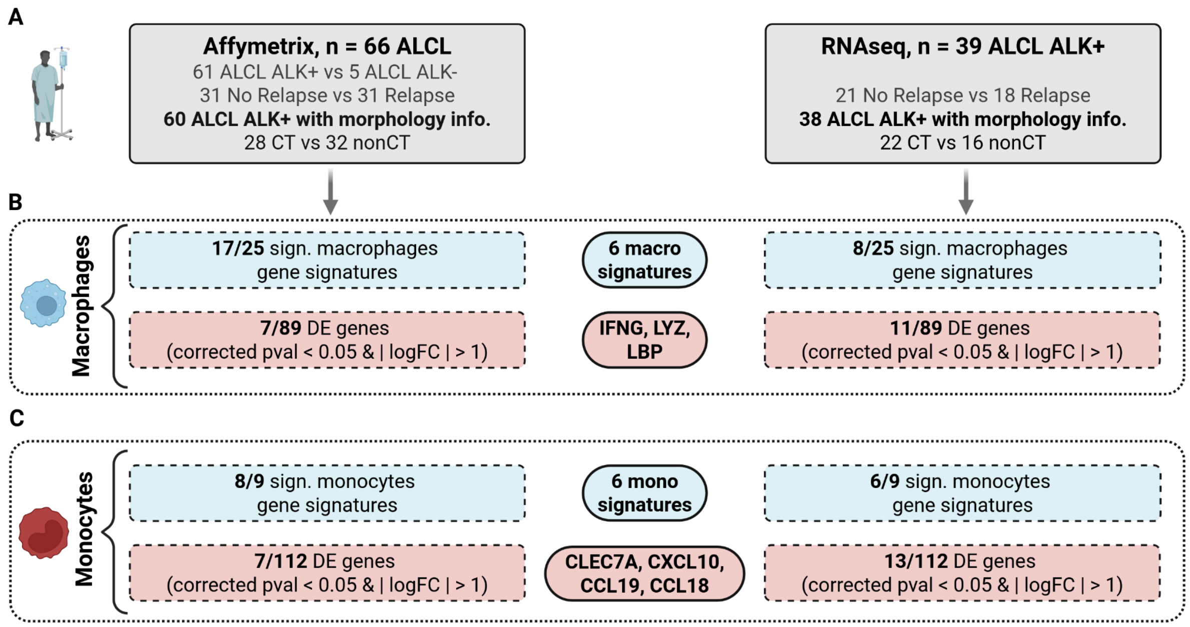

**Supplementary Figure S4: differential macrophages gene signatures between common (CT) and non-common-type (nonCT) variants. (A)** 17 signatures significatively different when comparing Sample Enrichment Scores (SES) between the 2 morphology groups on Affymetrix dataset (60 samples). **(B)** 8 signatures significatively different when comparing SES scores between the 2 morphology groups on the RNAseq dataset (38 samples).

Affymetrix and RNAseq common samples classified into the same enrichment clusters (cluster 1 and cluster 2) are in bold in both panels A and B. In the same way, common samples with discordant classifications are underlined. Macrophage gene signatures with significant discriminant scores in both Affymetrix and RNAseq are emphasised in bold.

**A. ALCL Affymetrix data**

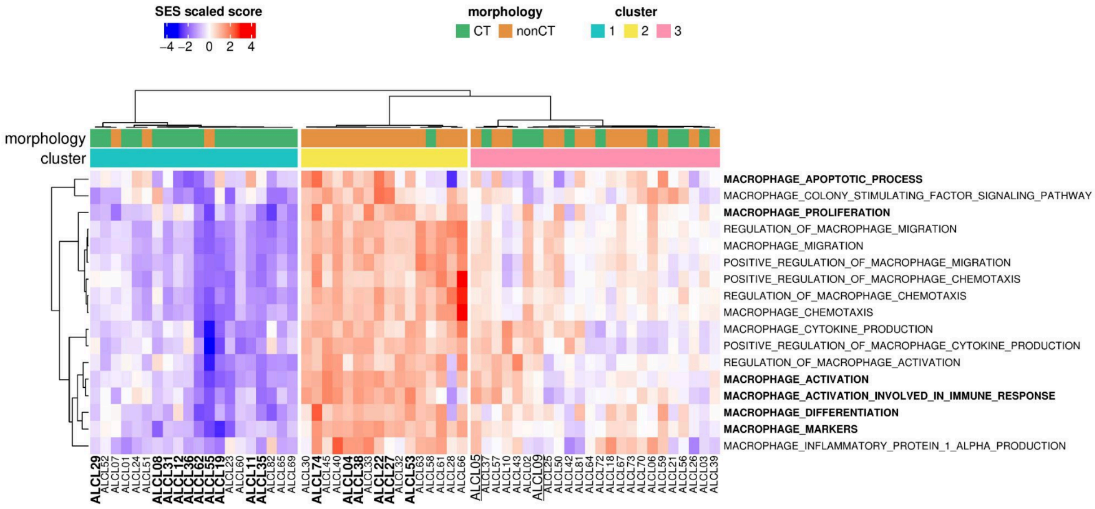


**B. ALCL RNAseq data**

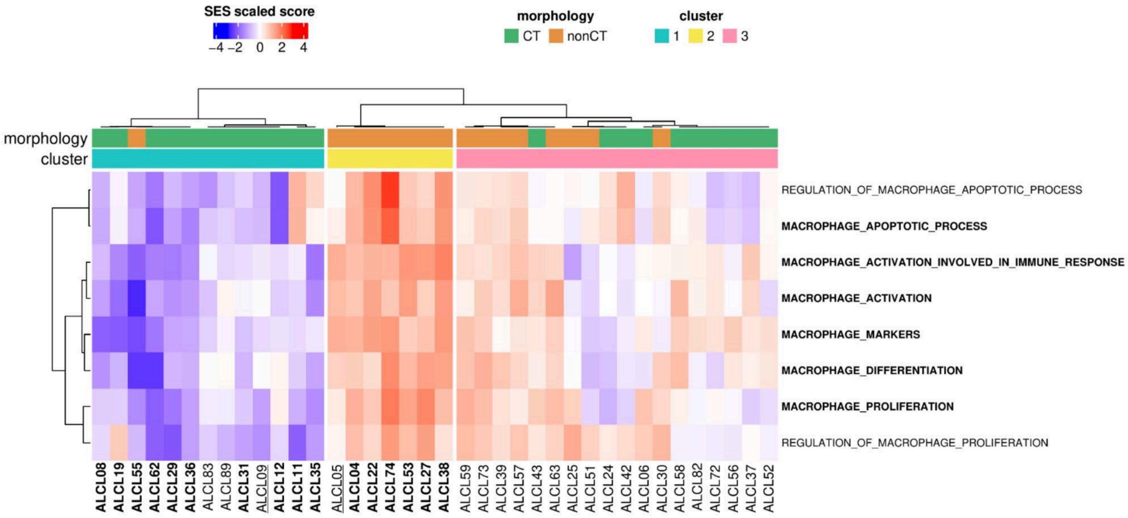

**Supplementary Figure S5: macrophage genes expression. (A)** 7 genes expression exploration into different normal T cells and ALK+ ALCL cell lines. Gene expression is normalized by the library sizes and in log10 scale. ZBTB46 is the only gene expressed mainly in sensitive or resistant to crizotinib ALK+ ALCL cell lines and PDX models. **(B)** Best transcriptomic signature to discriminate between common and non-common-type variants using Affymetrix data.

The genes both significatively differentially expressed within Affymetrix and RNAseq datasets are highlighted in bold.

**A. RNAseq** (CD3 T-lymphocytes and ALCL ALK+ cell lines or PDX)

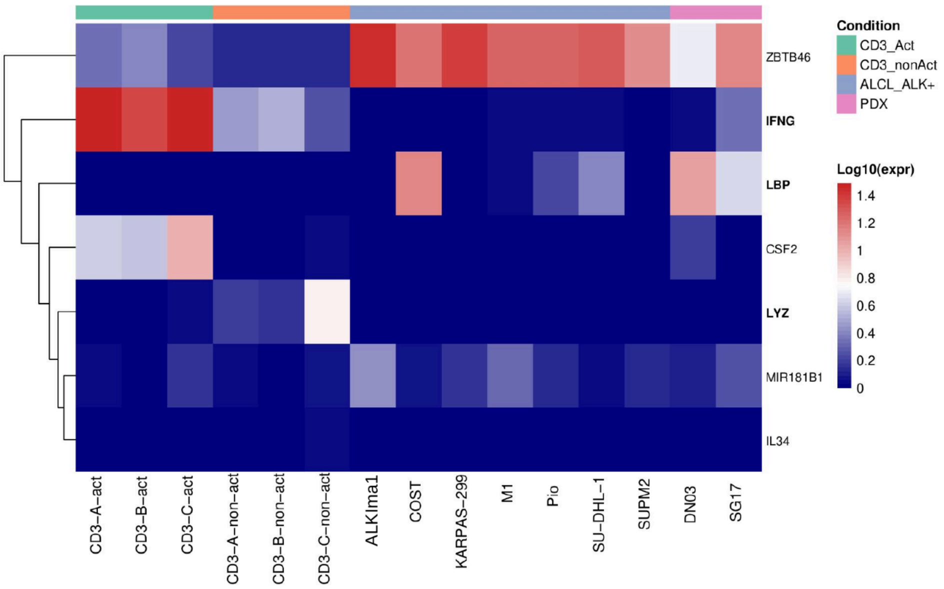


**B. Affymetrix**

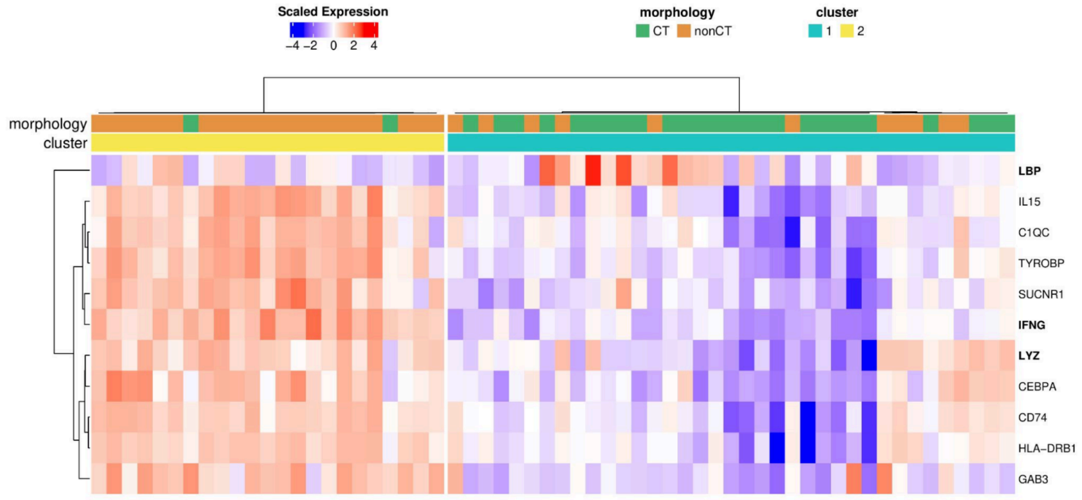

**Supplementary Figure S6: ALK-positive tumor cells and macrophages.** Small cell variant ALCL showing small ALK-positive tumor cells (ALK1 staining in yellow) mixed with a great number of histiocytes (CD68 staining in red).

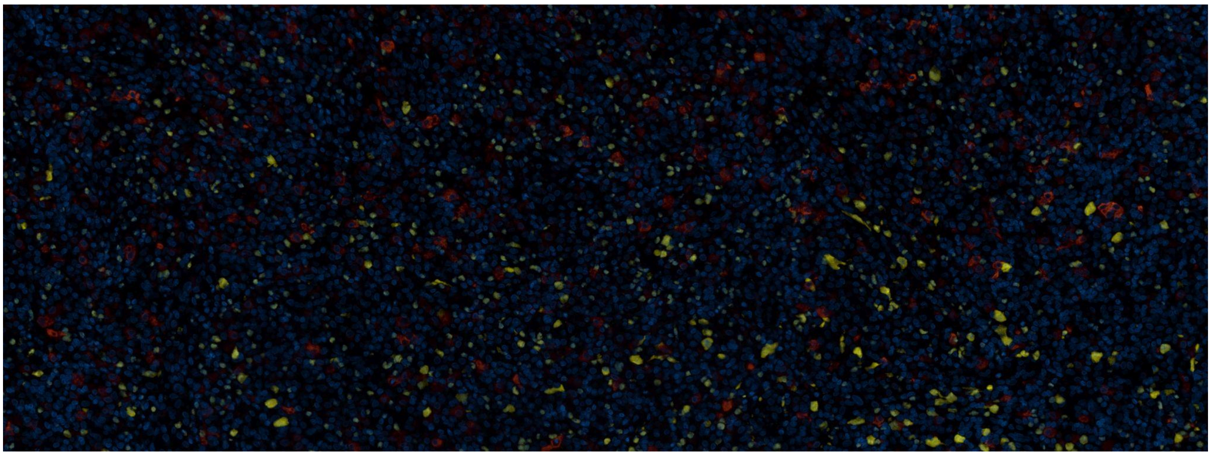

**Supplementary Figure S7: differential monocytes gene signatures between common (CT) and non-common-type (nonCT) variants. (A)** 8 signatures significatively different when comparing Sample Enrichment Scores (SES) between the 2 morphology groups on Affymetrix dataset (60 samples). **(B)** 6 signatures significatively different when comparing SES scores between the 2 morphology groups on the RNAseq dataset (38 samples).

Affymetrix and RNAseq common samples classified into the same enrichment clusters (cluster 1 and cluster 2) are in bold in both panels A and B. In the same way, the common sample with a discordant classification is underlined (ALCL82). Monocytes gene signatures with significant discriminant scores in both Affymetrix and RNAseq are emphasised in bold.

#### A. ALCL Affymetrix data

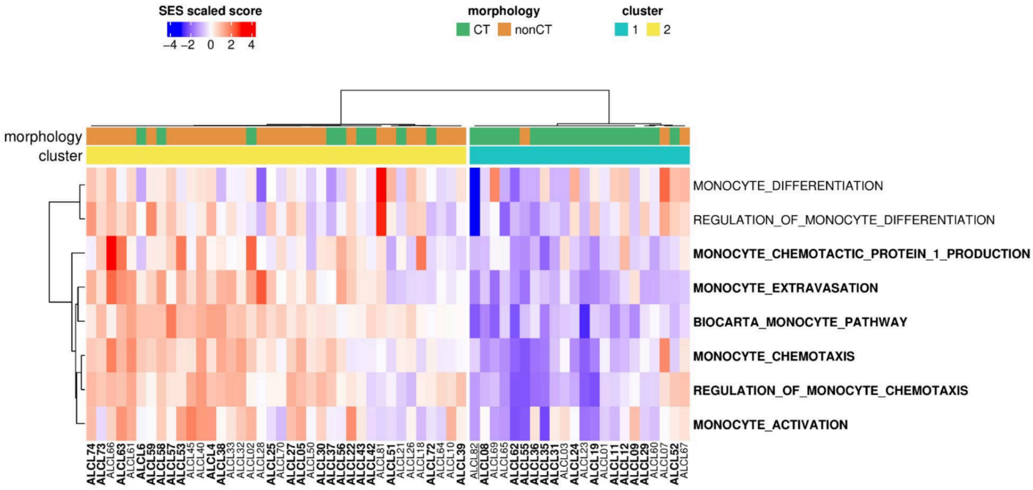


#### B. ALCL RNAseq data

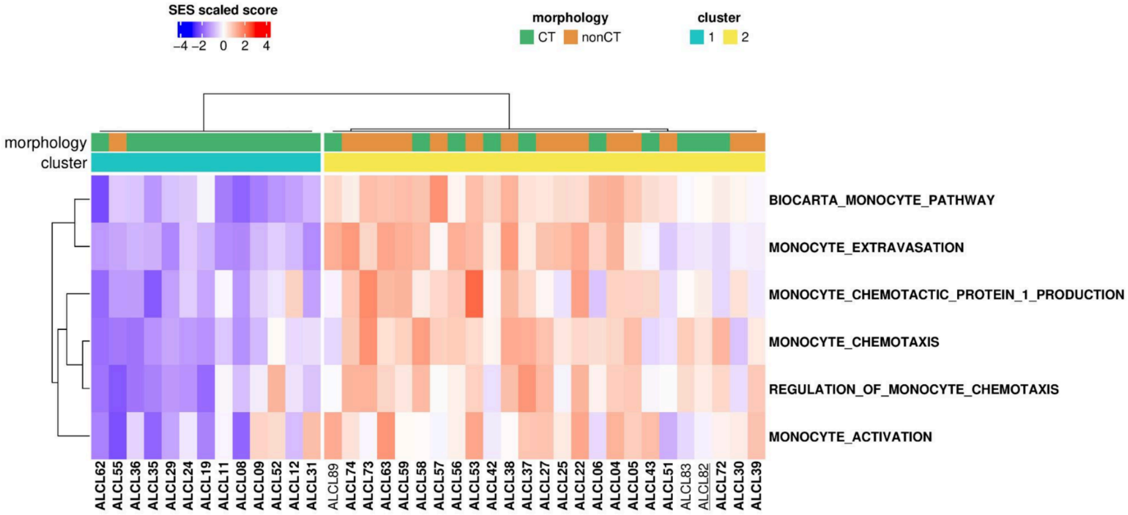

**Supplementary Figure S8: monocyte gene expression. (A)** 7 gene expression exploration into different normal T cells and ALK+ ALCL cell lines. Gene expression is normalized by the library sizes (see Methods) and in log10 scale. **(B)** Best transcriptomic signature to discriminate between common and non-common-type variants using Affymetrix data (corrected p-value < 0.05 and abs(logFC) > 1).

The genes both significatively differentially expressed within Affymetrix and RNAseq datasets are highlighted in bold.

**A.** **RNAseq** (CD3 T-lymphocytes and ALCL ALK+ cell lines or PDX)

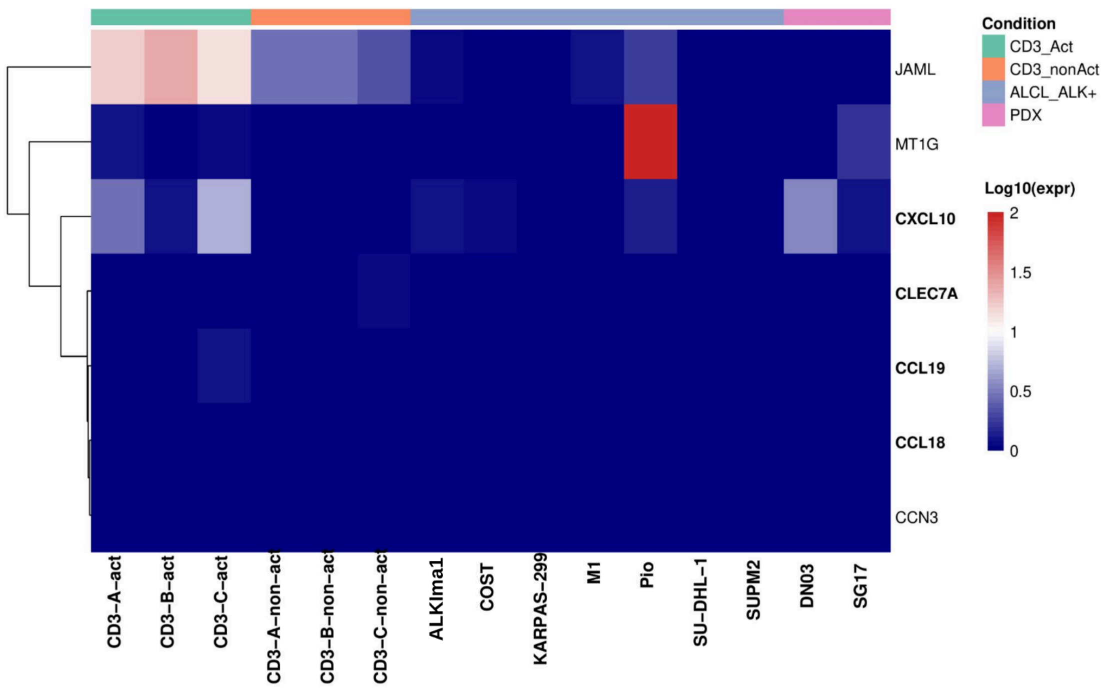


**B.** **Affymetrix**

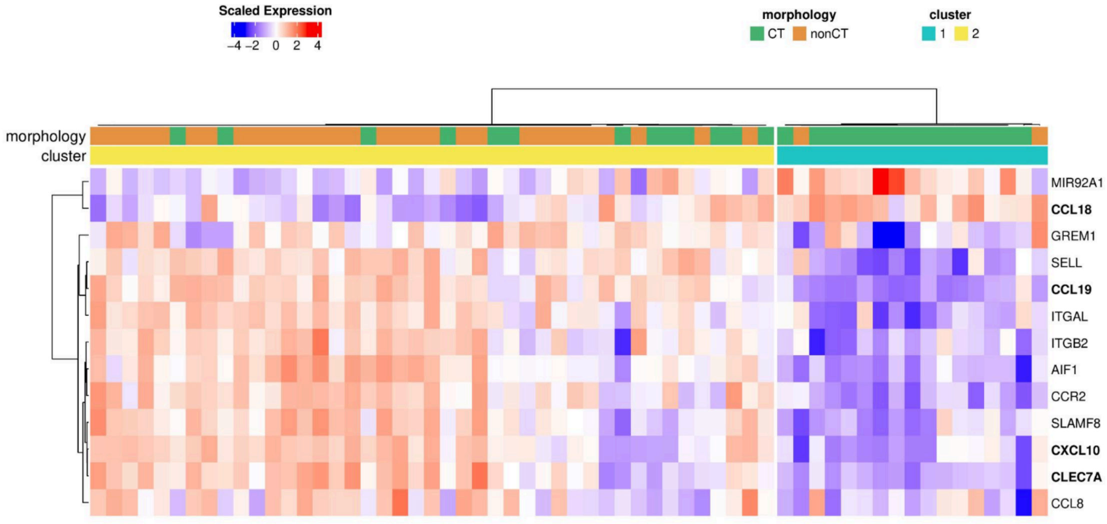
